# Neurobiological and Chemical Characterization of the Cyanobacterial Metabolite Veraguamide E

**DOI:** 10.1101/2025.06.19.660581

**Authors:** Jesus Sotelo, Sahar Mofidi Tabatabaei, Christian Fofie, Kelvin Fosu, Joseph Dodd-o, Rebecca Simcik, See Tack, Miguel J. Soto-Reyes, Saad Yousef, Eduardo J. Caro-Diaz, Vivek Kumar, Wade Van Horn, Benedict J. Kolber, Kevin J. Tidgewell

## Abstract

Ver E’s structure was validated by ¹H NMR, HRMS, and molecular networking analyses. Computational docking and NMR titration confirmed direct, saturable, and tight binding of Ver E to the human Sigma-2 receptor/transmembrane protein 97 (σ₂R/TMEM97). Functional calcium imaging in primary mouse sensory neurons revealed that Ver E increases intracellular Ca²⁺ levels without modulating store-operated calcium entry (SOCE). Multi-well microelectrode array experiments using human induced pluripotent stem cell (hiPSC) derived nociceptors showed that Ver E significantly reduced neuronal activity at physiological temperatures, but not under heat-stress conditions. Ver E exhibited no cytotoxicity at concentrations up to 30 µM in HEK293 cells, and immunocytochemistry confirmed that it does not alter phosphorylated eIF2α (p-eIF2α) expression, indicating a mechanism distinct from integrated stress response modulators. Collectively, these findings position Ver E as a non-toxic compound capable of selectively modulating neuronal excitability, thereby advancing the development of novel therapeutics for pain management.

**Significance:** Natural products have long been recognized as a rich source of therapeutics, accounting for over 60% of currently approved small-molecule drugs and underscoring their pivotal role in drug discovery. Marine cyanobacteria produce structurally diverse secondary metabolites with a wide array of biological activities. Among these are the veraguamides, a family of depsipeptides that have shown promise as future therapeutics in our recent studies. This work presents a detailed biological and chemical characterization of veraguamide E (Ver E), isolated from a Panamanian marine cyanobacterial collection. The σ₂R/TMEM97 system has been identified as a promising target to address unmet need for non-opioid therapeutics which can modulate neuronal excitability in the context of chronic pain. Discovery and identification of novel compounds which modulate this system can help us better understand its function as well as allow us to develop future therapeutics targeting this pathway.

**Highlights:**

- Veraguamide E specifically binds σ₂R/TMEM97 receptor with high affinity.
- Computational docking and NMR confirm a distinct binding mechanism.
- Ver E modulates calcium signaling in mouse DRG neurons and human iPSC-derived nociceptors.
- Ver E demonstrates no detectable cytotoxicity in human cell lines.

## INTRODUCTION

Millions of individuals worldwide suffer from chronic pain. Whether it presents as neuropathic pain, inflammatory pain, or postoperative pain, it is one of the most debilitating and pervasive health issues affecting communities and the leading reason individuals seek medical care^1^. Currently, a variety of pharmacological agents are available, including non-steroidal anti-inflammatory drugs (NSAIDs), opioids, and adjunctive therapies. However, these therapeutics can cause adverse effects, including addiction, respiratory depression, and death^2–4^. The limitations of existing treatments underscore the need for new, safer analgesics.

Marine organisms utilize specialized biosynthetic pathways to produce structurally diverse bioactive secondary metabolites for chemical communication and defense. These compounds have drawn significant scientific interest over the past two decades, leading to the exploration of their anticancer^5^, antibacterial^6^, antimalarial^7,8^, anti-inflammatory^9^, and neuroactive^10,11^ properties.

Within this spectrum of bioactive compounds are the veraguamides, which are a specific type of lipopeptides called depsipeptides. Their unique defining features include a conserved proline residue, multiple N-methylated amino acids, α-hydroxy acid, and a C8-polyketide-derived *β*-hydroxy acid moiety, which has characteristic terminal groups that vary, appearing as an alkynyl bromide, alkyne, or vinyl group. These depsipeptides were originally discovered from cyanobacterial collections^12,13^, and have since been the focus of increased biochemical and pharmacological exploration. In this work, we examine Ver E which phylogenetic analysis suggests is produced by an *Okeania sp*^14^ and is a representative of the terminal alkyne containing cyclic peptides creating strong bioactivity. In this research, we tested two elements of Ver E’s chemistry and biology. First, we evaluated the potential of Ver E to bind to an emerging new target for analgesic drug development, the sigma-2 receptor/transmembrane protein 97 (σ₂R/TMEM97)^15,16^. This non-canonical transmembrane protein is associated with the endoplasmic reticulum and has been implicated in a diverse array of cellular functions including cholesterol and calcium homeostasis^17–22^. In the context of pain, modulators of σ₂R/TMEM97 reduce neuropathic pain^23,24^ for extended periods of time that exist beyond the predicted half-life of the compounds. σ₂R/TMEM97 is highly expressed in sensory neurons of the dorsal root ganglion of mice and humans, suggesting that the analgesic effects are mediated, at least in part, peripherally^23,24^. This pharmacological profile makes σ₂R/TMEM97 an exciting target of exploration for pain therapeutics. Recently, we found that a cousin of Ver E, veraguamide O, showed affinity for σ₂R/TMEM97 suggesting that other members of the veraguamide family may bind the protein^14^

Second, related to the potential for Ver E to bind a protein known to be in sensory neurons, we sought to explore the impact of Ver E on sensory neuron function. We explored calcium signaling of primary mouse dorsal root ganglia (DRG) neurons during Ver E treatment. Calcium signaling is an important part of sensory neuron signaling that is known to be (1) modulated by σ₂R/TMEM97 and (2) critical to nociceptor function and pain^25,26^. Studies have demonstrated that modulations in calcium influx or SOCE can either potentiate or attenuate pain signals^27^. Other marine lipopeptides such as barbamide have been reported to act on calcium channels and influence pathways relevant to neuronal excitability^10^. Consequently, testing the impact of Ver E on calcium dynamics in DRG neurons is a promising approach for investigating this compound’s analgesic potential. Next, we evaluated the impact of Ver E on sensory neuron excitability using human induced pluripotent stem cell (hiPSC) derived nociceptors, an assay with improved translational relevance^28^. Finally, we also tested the hypothesis that specificity for Ver E’s effects by σ₂R/TMEM97 may lead to modulation of the integrated stress response (ISR). The ISR is caused by ER stress and is known to be associated with chronic pain^29,30^ and σ₂R/TMEM97 modulation^23^.

Overall, we found that Ver E is a structurally distinct compound compared to other identified σ₂R/TMEM97 ligands and from NMR binding and hIPSC studies clearly has potential for functional activity in humans. Ver E provides several points of modification, making it a rich scaffold for therapeutic development and tool compound generation. Moreover, Ver E lacks cytotoxicity and modulates both mouse and human nociceptors consistent with future translational utility. These data suggest that Ver E may mediate its biological effects through σ₂R/TMEM97 but lacks an effect on the ISR providing a unique cellular mechanism beyond those of other recently published σ₂R/TMEM97 modulators^15,17–20,23,24^. This combined utilitarian chemical structure and biological profile provide impetus for continued exploration of this and related scaffolds for further development.

## RESULTS

### Collection, Isolation, and Identification of Ver E

A cyanobacterial sample (DUQ0008) identified as *Okeania sp.* was collected from the coastal waters of Isla Mina in the Las Perlas Archipelago, Panama (GPS coordinates: N 8°29.717′, W 78°59.947′)^14^. From this collection, 2.8 mg of a colorless oily compound was purified from a fraction eluting with 70% methanol / 30% ethyl acetate using multiple rounds of chromatography (**Supplemental Figures S1, S2**).

^1^H NMR and COSY spectra of the purified compound were highly consistent with veraguamide depsipeptides previously isolated from the same collection (**Figure 1**)^14^. By comparing the spectra with literature reports, the compound was identified as Ver E (**1**)^12,13^. Diagnostic NMR signals include nine methyl groups between 0.85–1.23 ppm, two sharp N-Me singlets at 2.92 ppm and 2.99 ppm, an acetylene CH at 3.14 ppm, proline diastereotopic hydrogens at 3.60 ppm and 3.83 ppm, five deshielded CHs at 4.04, 4.21, 4.72, 4.82, and 4.91 ppm, and an NH peak at 6.28 ppm^13^ (**Figure S3, Figure S4**). HRMS (positive mode) confirmed the molecular formula C_39_H_64_N_4_O_8_, showing peaks at m/z 717.4766 [M + H]^+^ and 739.4581 [M + Na]^+^ (calculated for C_39_H_65_N_4_O_8_, 717.4802) (**Figure S5**). Utilization of MS/MS classical molecular networking through the GNPS2 platform of the pure compound also provided strong evidence for it to be a veraguamide and specifically Ver E. The MS/MS fingerprint for the compound yielded a cosine similarity score of 0.4 compared to the Ver E MS/MS fragmentation pattern, with 15 matching peaks identified^31,32^ (**Figure S6**). The compound networked with two other veraguamide analogs (**Figure 1**) with minor but distinct modifications of amino acid side chains, in the case of veraguamide D, and in amino acid side chains and HMOYA modification for veraguamide G. Interestingly there is also connection into a network containing Dudawalamide A^33^, another marine cyanobacterial cyclic depsipeptide with a DHOYA rather than HMOYA lipid tail. Based on HR-MS, key diagnostic NMR signals, and the molecular networking analysis we confidently identify the compound as Ver E.

**Figure 1:**
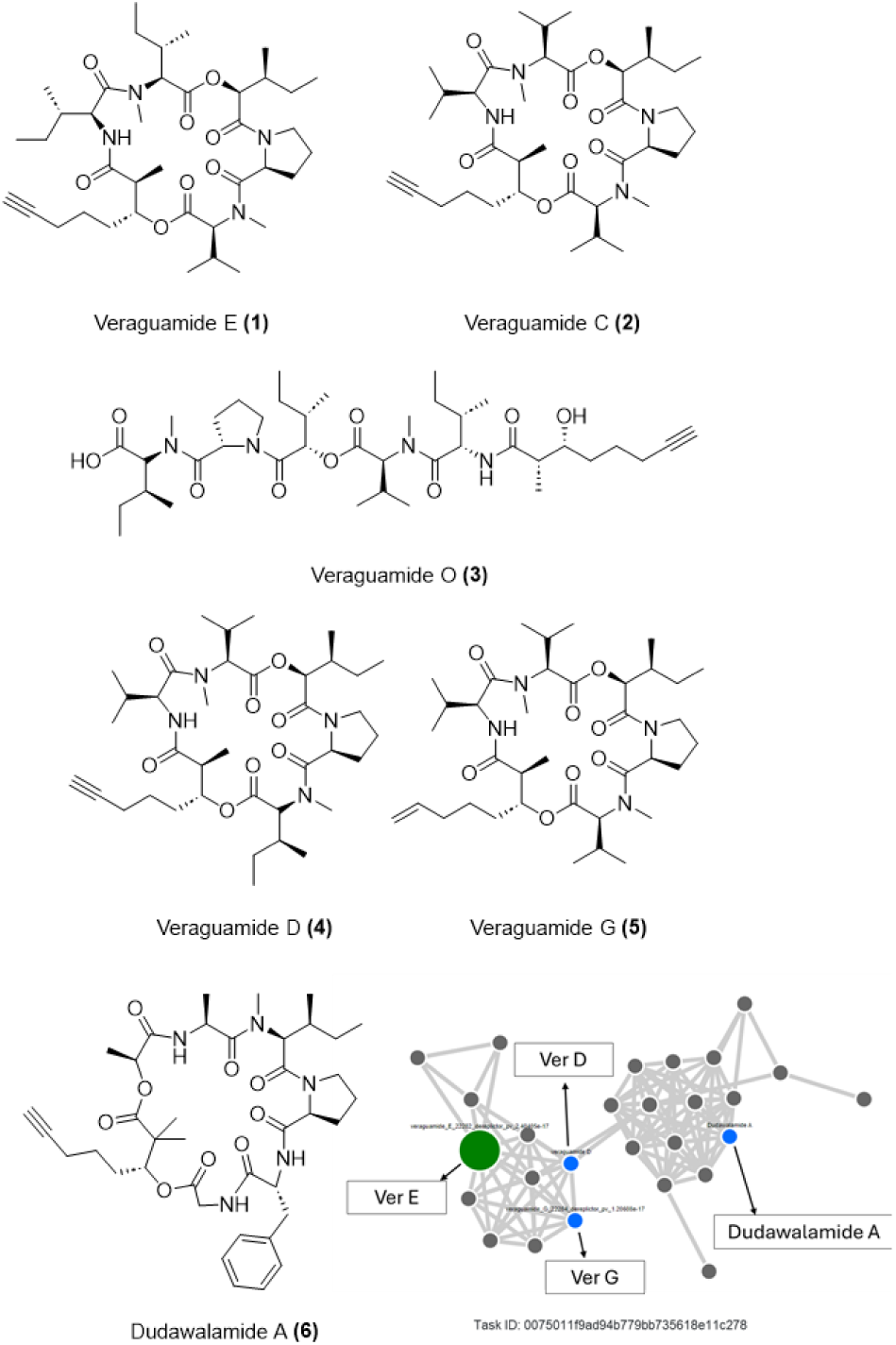
Chemical Structures and Molecular Network. Chemical structures of veraguamide E (**1**), veraguamide C (**2**), veraguamide O (**3**), veraguamide D (**4**), veraguamide G (**5**), dudawalamide A (**6**), and GNPS2 Molecular Network of VerE task ID: 0075011f9ad94b779bb735618e11c278.

### Ver E Occupies a Distinct Chemical Space among σ₂R/TMEM97 Ligands

To better understand how Ver E compares to known σ₂R/TMEM97 specific ligands, we performed similarity calculations between Ver E and a select group of previously described σ₂R/TMEM97 ligands^15^ using an Overlap Analysis tool (InstantJChem, ChemAxon). The computational comparison produced Tanimoto scores that quantify the structural similarity of Ver E to each σ₂R ligand selected (**Figure 2**)^34^. Ver E holds the highest similarity with SM-21 and histatin-1 (entries 4 and 17, **Table S1**), with a median of 0.32 across all σ₂R/TMEM97 ligands, suggesting that Ver E is structurally distinct to established σ₂R/TMEM97 ligands.

**Figure 2:**
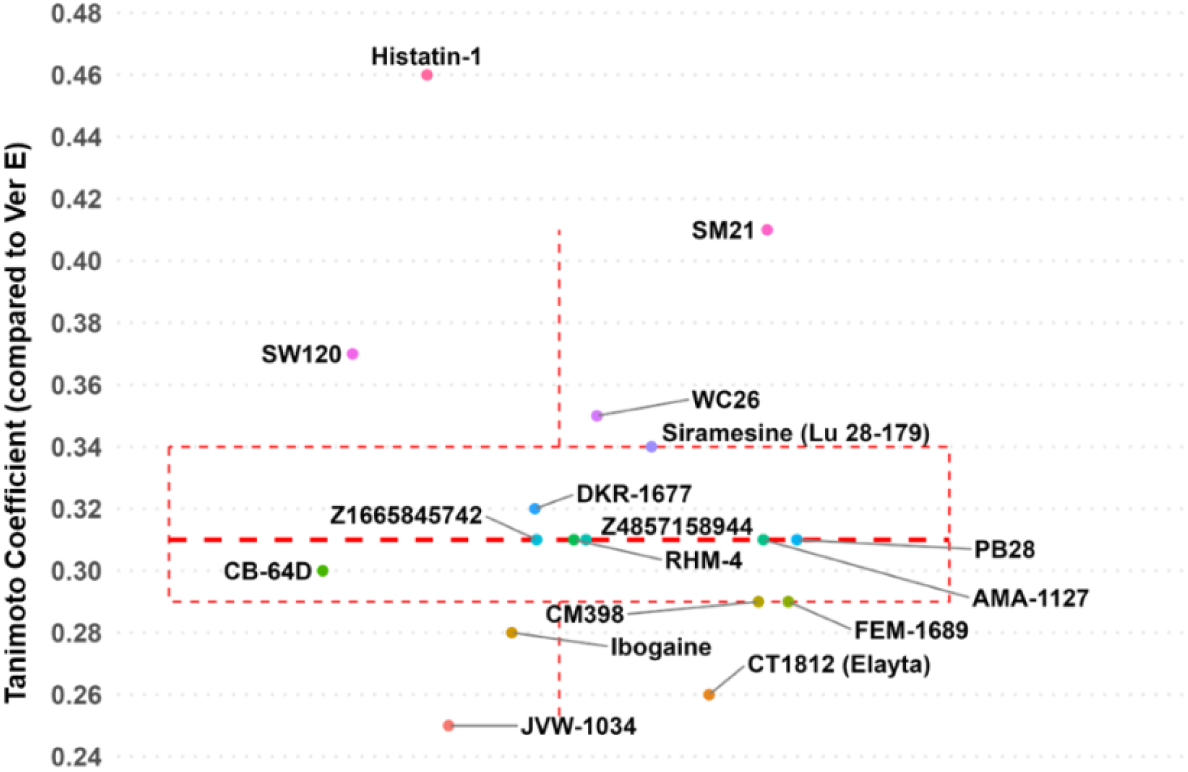
Box plot of Tanimoto Coefficients for σ₂R ligands compared to Ver E. Interquartile Range (IQR) of all coefficients is represented as a dotted red rectangle and red dashed lines representing Q1 (25^th^ percentile), median, and Q3 (75^th^ percentile).

To further compare Ver E to known σ₂R/TMEM97 ligands, we conducted predictions of physicochemical descriptors of Ver E and σ₂R/TMEM97 ligands using the DataWarrior platform^35^ (**Table S2**) and performed Principal Component Analysis (PCA) (**Figure 3, Figure S7**), where Ver E describes its own chemical space. Thus, comparative multivariate analysis of predicted physiochemical descriptors together with Overlap Analysis strongly supports the notion that Ver E represents a structurally unique σ₂R/TMEM97 ligand chemotype.

**Figure 3:**
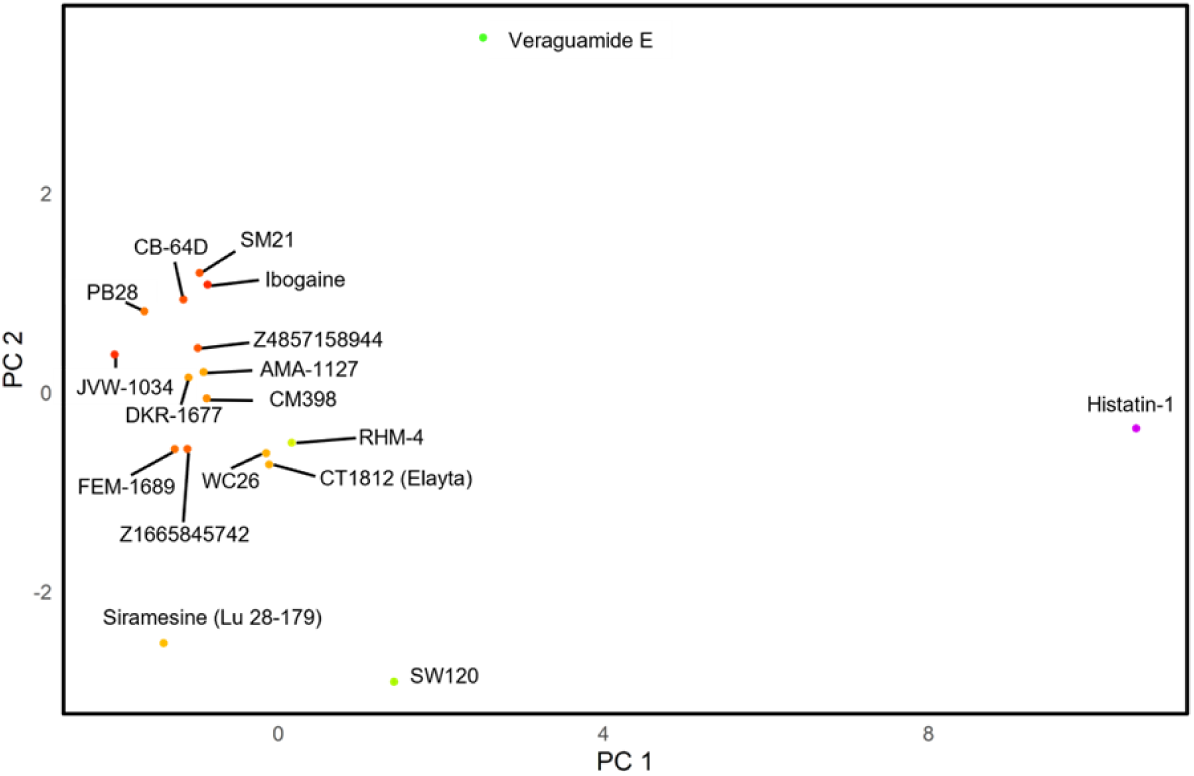
Principal Component Analysis (PCA) plot of 13 predicted physicochemical descriptors of Ver E and previously described σ₂R/TMEM97 ligands.

### Ver E Directly Binds σ₂R/TMEM97 at a Non-canonical Surface Site

Due to the unique structure of Ver E and our recently published data showing significant radioligand competitive binding of veraguamides to σ₂R/TMEM97^14,23,24,36^, we wanted to test Ver E’s potential molecular binding partners. In our previous report, both the cyclic and linear depsipeptides veraguamide C and O showed significant binding affinity to σ2/TMEM97 as screened by the Psychoactive Drug Screening Program (PDSP)^37^. While we were unable to test Ver E in this assay due to compound availability, we were able to experimentally determine its binding potential in σ₂R/TMEM97 using computational docking and protein NMR-detected titration experiments.

AlphaFold3 was employed to model the full-length σ₂R/TMEM97 receptor^38^ (**Figure 4**). Consistent with previously published modeling exercises, the σ₂R/TMEM97 was predicted to conform to the same architecture as the published bovine σ₂R/TMEM97 receptor^39^. We employed both generative diffusion-based and energy-based methods to predict the binding pocket (**Figure S8**). Consistent with the protoypical σ₂R/TMEM97 PB28^39^, the predicted binding pocket is in the concave region of the σ₂R/TMEM97. Unlike PB28, however, we have found that Ver E is likely to reside outside of the recessed region of the receptor (**Figure 4E**, **Figure 4H**). Energy of binding as assessed using AutoDock-GPU predicts binding energy similar to existing ligands of σ₂R/TMEM97.

**Figure 4:**
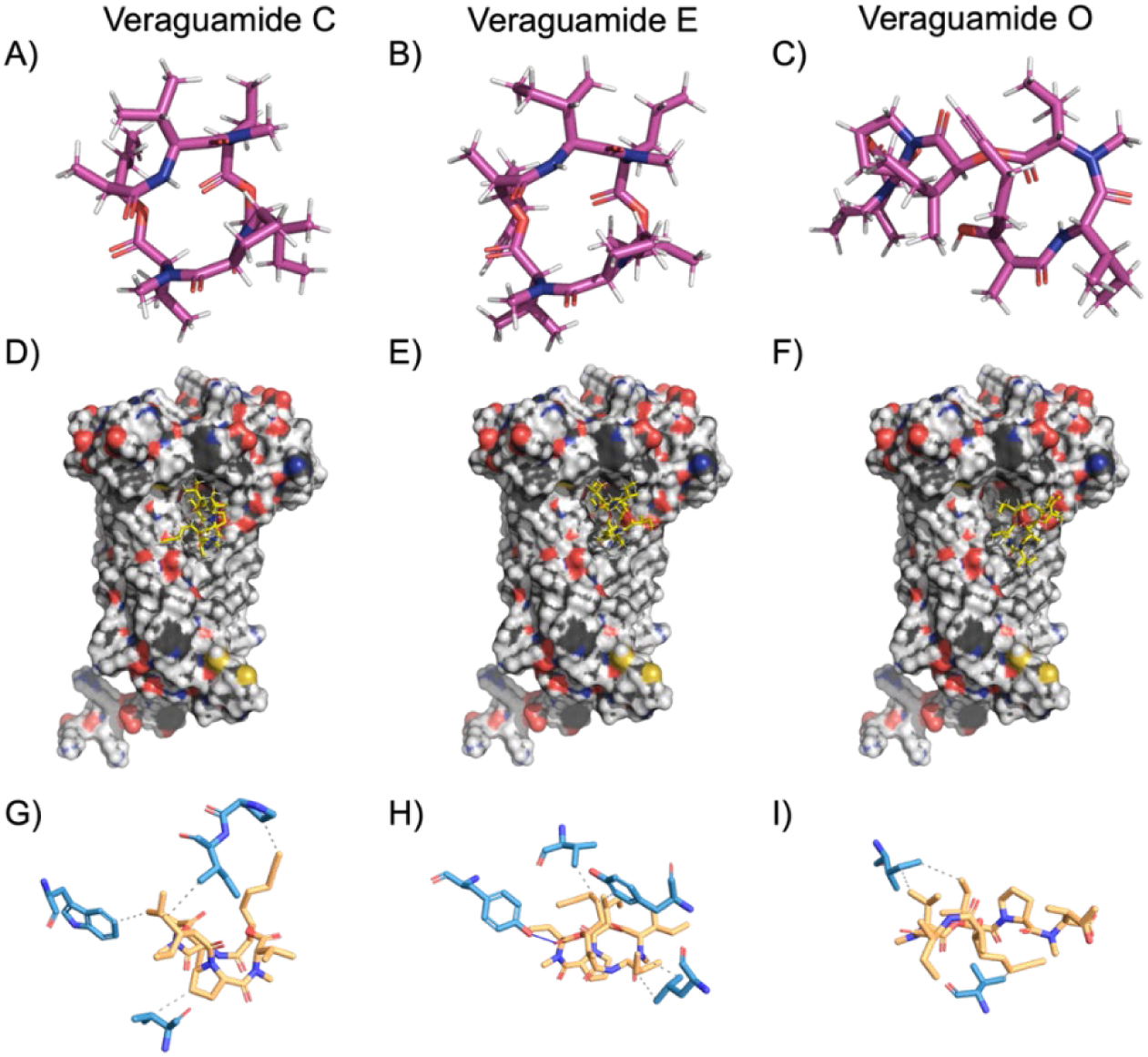
*In silico* prediction of binding pocket. **(A-C):** 3D representations of the ligand conformers, generated using obgen in a Universal Force Field. **(D-F)** Most energetically favorable binding conformation against AlphaFold3 generated the σ₂ receptor. **(G-I)** Protein-Ligand Interaction Profiles of the ligand’s predicted binding pocket.

Next, we tested the potential for direct interactions between Ver E and reconstituted human σ₂R/TMEM97 (**Figure S9**) using NMR-detected σ₂R-ΔER binding studies. ^1^H-^15^N BEST-TROSY-NMR experiments were recorded at increasing Ver E concentrations. Global evaluation of the binding data by principal component analysis and discrete local chemical shift perturbation binding analysis showed saturable binding isotherms, indicating direct and specific Ver E binding to σ₂R/TMEM97 (**Figure 5**, **S10**). The binding data were fit to a single-site binding model, yielding a global *K*_d_ of 0.3 ± 0.2 nmol%, confirming that σ₂R/TMEM97 in DPC micelles directly binds Ver E. A control binding experiment was performed, mimicking the vehicle concentrations. This control titration showed no non-specific binding to σ₂R/TMEM97 (**S11**), validating a direct interaction between Ver E and σ₂R/TMEM97.

**Figure 5:**
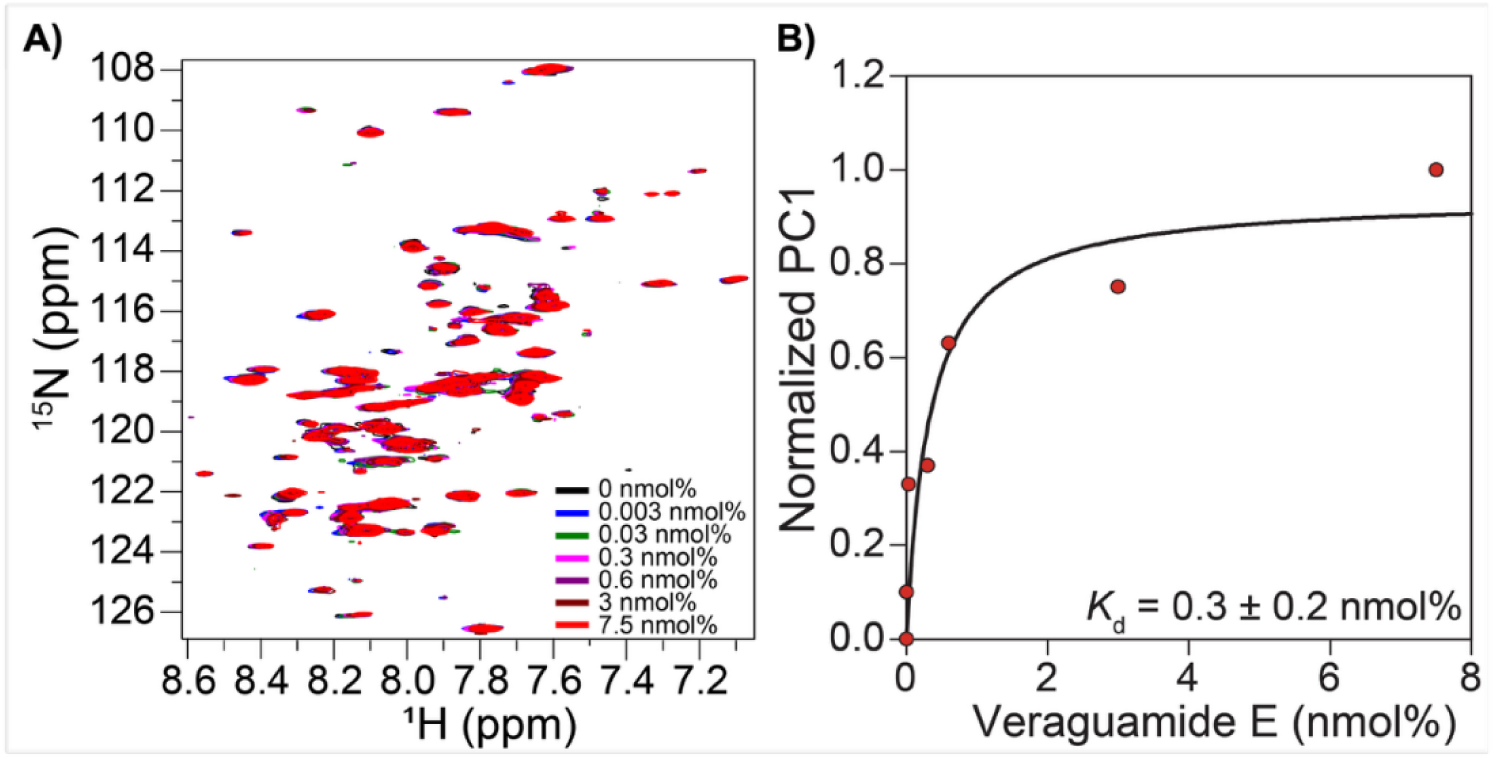
Human σ₂R/TMEM97 directly binds Ver E. **A)** Superimposed ^1^H-^15^N BEST-TROSY spectra of σ₂R/TMEM97 titration with Ver E at 37 °C. **B)** Nonlinear fitting of chemical shift perturbations to a single-site binding model reveals a global *K*_d_ of 0.3 ± 0.2 nmol% for Ver E binding to σ₂R/TMEM97.

### Ver E Elevates Intracellular Ca^2+^ without Modifying Store-Operated Entry in DRG Cells

Next, we tested the impact of Ver E in several biological assays, starting with measuring calcium (Ca^2+^) levels in mouse DRG *in vitro*. We investigated the potential of Ver E to modulate intracellular Ca²⁺ levels and, by association, alter nociceptive neuronal signaling. Since marine-derived depsipeptides have been shown to influence neuronal signaling by targeting calcium channels and pathways^40^, we hypothesized that Ver E may similarly affect Ca²⁺ dynamics in DRG neurons. To test this, we performed functional calcium imaging recordings on DRG from mice expressing the genetically encoded calcium indicator GCaMP6f. We measured both baseline calcium flux and the ability of Ver E to directly influence DRG excitability under various treatment conditions, including its effects on SOCE, a key pathway for maintaining intracellular calcium homeostasis^41,42^.

To assess its impact on calcium homeostasis, 10 µM Ver E was applied directly to DRG neurons, and changes in calcium signaling were monitored during a SOCE activation protocol (**Figure 6**). In this experiment, the Sarcoendoplasmic Reticulum Calcium ATPase (SERCA) pump inhibitor thapsigargin was applied with or without Ver E in the absence of extracellular Ca²⁺. Under these conditions, thapsigargin induces the release of intracellular Ca²⁺ stores. Upon reintroduction of extracellular Ca²⁺, the SOCE response was measured. This approach enables evaluation of both SOCE buildup during thapsigargin treatment and the subsequent Ca²⁺ influx associated with homeostatic recovery. Ver E significantly increased DRG excitability during its co-application with thapsigargin compared to thapsigargin alone (P < 0.01), indicating an acute effect on intracellular Ca²⁺ levels. However, no significant change was observed in the subsequent SOCE response, suggesting that Ver E’s mechanism of action does not directly target store-operated calcium pathways.

**Figure 6.**
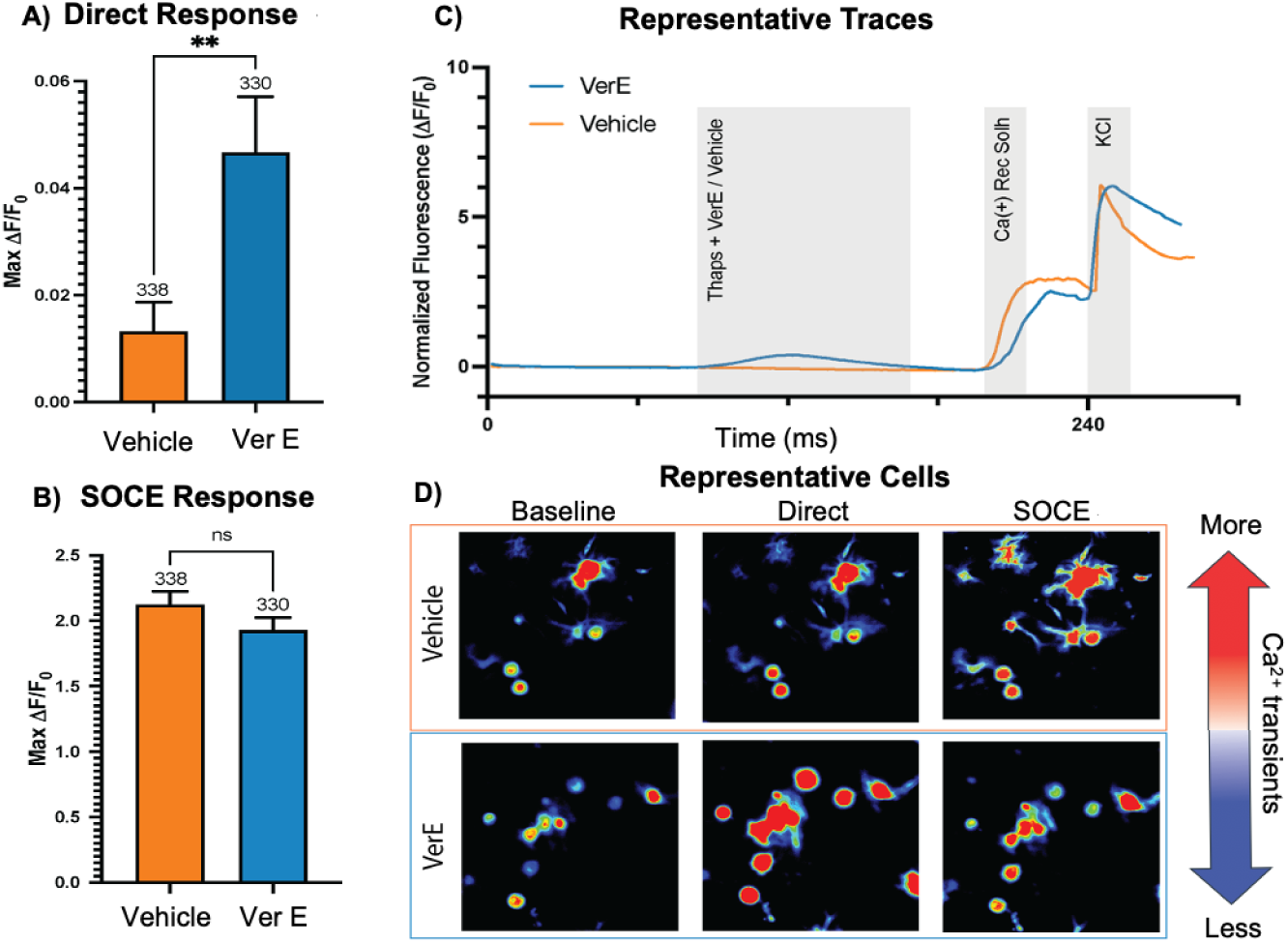
Effect of Ver E on Calcium Homeostasis in DRG Neurons. **(A)** Maximum fluorescence measured in DRGs exposed to vehicle (0.1% DMSO) or Ver E (10 µM). Ver E treatment significantly increased fluorescence compared to vehicle (unpaired two-tailed t-test, \*\**P* < 0.01). Error bars recorded as standard error of mean (SEM) with n (number of cells) indicated in graph. **(B)** Maximum fluorescence observed during the SOCE response with vehicle or Ver E. No significant difference was detected between groups. **(C)** Representative real-time calcium imaging traces for vehicle vs. Ver E-treated DRGs. Gray bars indicate the period of exposure to each treatment. **(D)** Representative heat map images of DRG cells illustrating the direct calcium response (left) and the SOCE response (right) under vehicle or Ver E treatment.

### Ver E Transiently Dampens Spontaneous Nociceptor Firing without Affecting Heat-Evoked TRPV1 Activity

DRG neurons also play a key role in peripheral nociception and are known to exhibit both spontaneous and evoked firing. To model this activity *in vitro*, we utilized hiPSC-derived nociceptors cultured on multi-electrode array (MEA) plates. By day 28, these neurons show spontaneous firing, reflecting baseline hyperexcitability characteristic of neuropathic pain conditions^28^. Application of Ver E at 30 µM significantly reduced neuronal activity by approximately 60% (P < 0.05) within the first hour at 37 °C; however, this inhibitory effect was transient, with activity returning to baseline by the 24-hour time point (**Figure 7**). In contrast, during transient heat ramp applications from 37 °C to 42 °C, Ver E did not produce a reduction in activity compared to vehicle-treated cells. These results indicate that Ver E does not block TRPV1 channels which normally open in response to noxious heat to depolarize neurons. In turn, that initial depolarization recruits voltage-gated sodium (Naᵥ), potassium (Kᵥ), and calcium (Caᵥ) channels, which together drive action potential firing in sensory neurons exposed to high temperatures.

**Figure 7:**
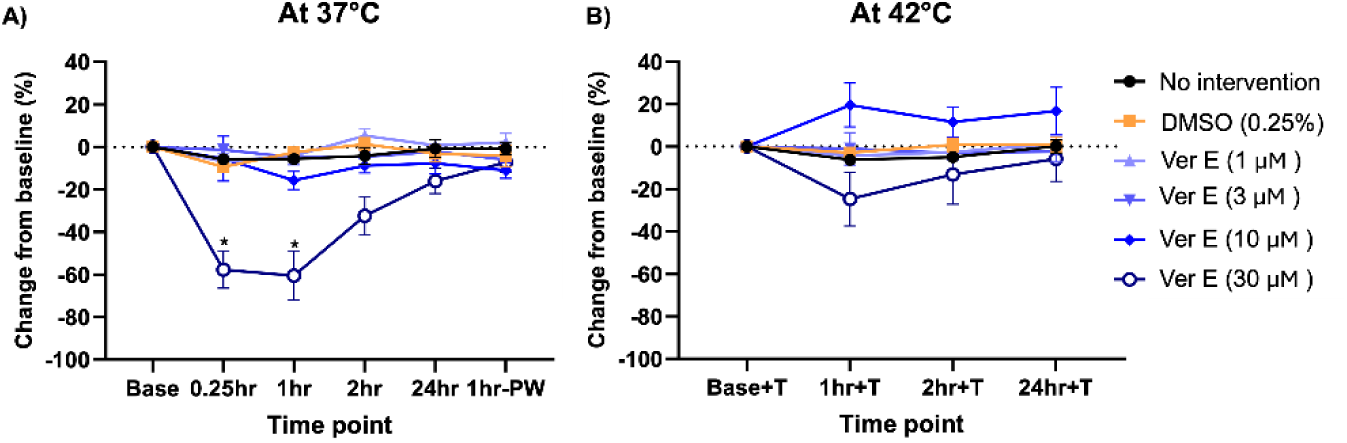
Inhibitory effects of Ver E on sensory neuron activity *in vitro*. **A)** Ver E significantly reduced the activity of hiPSC-derived nociceptors at a concentration of 30 µM at 37°C. **B)** A non-significant decrease was observed at 42°C. Data are presented as Mean ± SEM for each concentration point. *p < 0.05 indicates a significant difference compared to DMSO-treated cells at the indicated time point, as determined by two-way ANOVA with Dunnett’s post hoc test.

### Ver E Does Not Alter p-eIF2α Levels, Leaving the Integrated Stress Response Intact in DRG Neurons

Next, we investigated whether Ver E modulates the integrated stress response (ISR) in mouse DRG neurons. The ISR is a conserved cellular pathway that regulates protein synthesis in response to stress and has been implicated in pain sensitization and neurodegenerative disease^21,23,29^. Prior work has shown that certain σ₂R/TMEM97 modulators can disrupt this pathway by reducing phosphorylation of the key ISR regulator eIF2α^23^. To assess whether Ver E affects ISR signaling, we performed immunocytochemistry for p-eIF2α in DRG neurons treated with 100 nM or 1000 nM Ver E. No observable differences in p-eIF2α levels were detected compared to vehicle-treated controls (**Figure 8**). In contrast, treatment with FEM 1689, a known σ₂R/TMEM97 modulator produced a marked reduction in p-eIF2α expression at 30 nM and 100 nM, consistent with published data^23^. Neuronal identity was confirmed by co-staining for peripherin. These results suggest that Ver E does not affect ISR signaling through eIF2α phosphorylation under these conditions.

**Figure 8.**
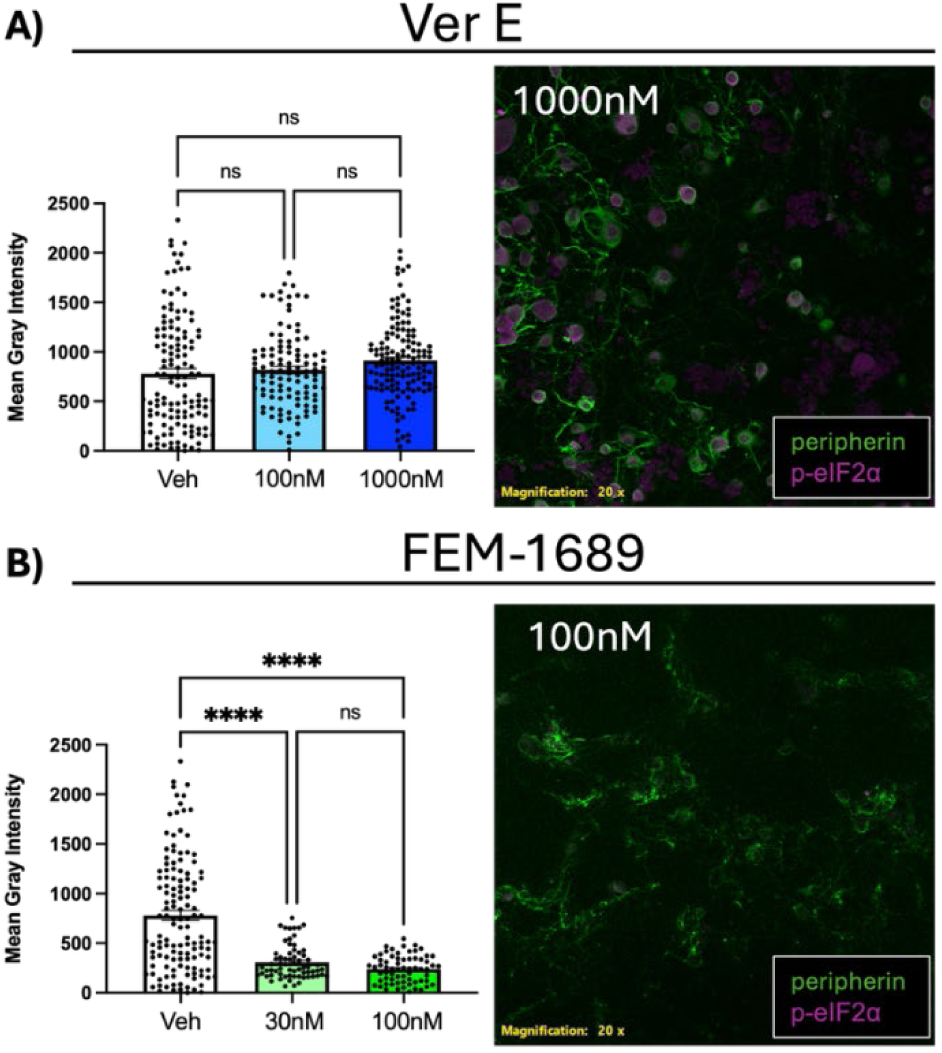
Impact of Ver E on the Integrated Stress Response. **A)** Effect of Ver E (100 nM and 1,000 nM) on p-eIF2α expression in primary mouse DRGs. No significant difference was observed compared to vehicle. Representative immunocytochemistry image on right of primary mouse DRGs treated with Ver E (1000 nM), visualized for p-EIF2α (magenta), and peripherin (green). **B)** Effect of FEM 1689 (30 nM and 100 nM) on p-eIF2α expression in primary mouse DRGs. FEM 1689 significantly decreased the mean gray intensity compared to vehicle. **C)** Representative immunocytochemistry image on right of primary mouse DRGs treated with FEM-1689 (100 nM), visualized for p-EIF2α (magenta), and peripherin (green). Data are presented as mean ± SEM. Statistical significance was determined by one-way ANOVA with Tukey’s multiple comparisons (****P < 0.0001).

### Ver E Shows No Detectable Cytotoxicity in HEK293 Cells up to 30 µM

Next, we assessed the potential cytotoxicity of Ver E. In parallel with its potential impact on neuronal calcium signaling, Ver E’s therapeutic viability also depends on its safety and selectivity. Marine natural products, though often pharmacologically potent, can also exhibit cytotoxicity that limits their therapeutic potential^12^. Early assessment of cytotoxicity in mammalian cell lines such as HEK293 cells is critical to identify compounds with acceptable safety profiles. HEK293 cells, derived from human embryonic kidney tissue, are widely used because they are easy to culture, genetically stable, and represent a relevant human cellular context for initial toxicity screening. Evaluating Ver E’s cytotoxic profile in this system not only informs safe dosing ranges but also aids in prioritizing candidate molecules for further development and optimization. Previous studies have reported that certain veraguamides exhibit varying degrees of cytotoxicity^12^. However, we recently showed that veraguamide C was non-toxic to HEK293 cells and exhibited only mild cytotoxicity in breast cancer cell lines^14^. To evaluate Ver E, we tested concentrations ranging from 0.03 µM to 30 µM using HEK293 cells, with digitonin included as a positive cytotoxicity control. Ver E showed no evidence of cytotoxicity across the tested concentration range (**Figure 9**). In contrast, digitonin induced a clear cytotoxic response, with an LC₅₀ of 9.68 µM and complete cell death observed at 30 µM. These findings indicate that the LC₅₀ for Ver E exceeds 30 µM under these conditions, supporting its safety profile and distinguishing it from more toxic analogs.

**Figure 9:**
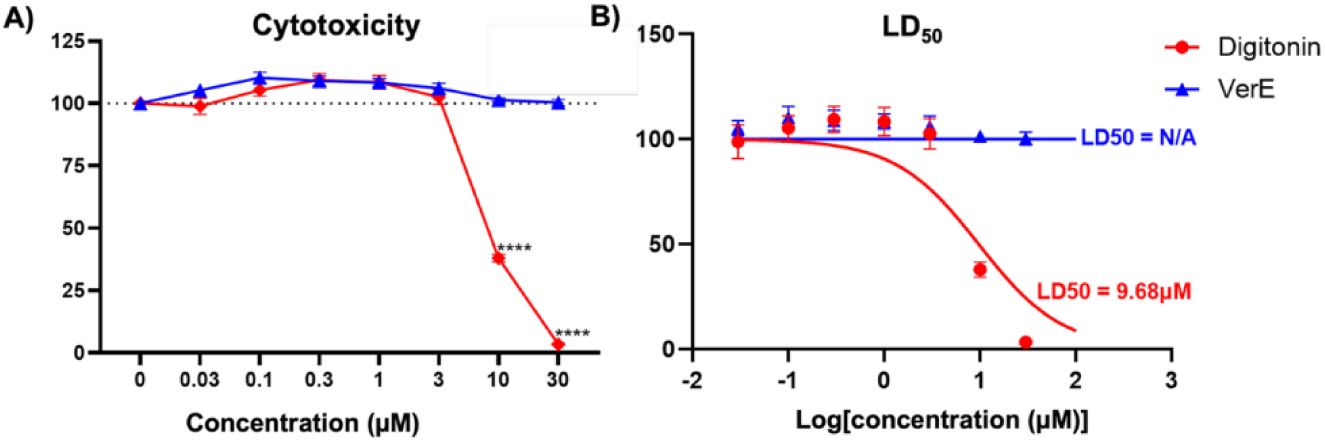
Cytotoxicity profile of Ver E compared to digitonin across a concentration range of 0.03–30 µM. **A)** At 30 µM, digitonin caused 100% cell death, whereas Ver E showed no cytotoxicity at the same concentration. **B)** Based on the logarithmic non-linear regression curve, the LD_50_ of digitonin was determined to be 9.68 µM, while Ver E had no detectable LD_50_, as it did not induce cell death at any tested concentration. **A) & B)** Each point represents a Mean±SEM. ****p<0.0001 indicates a significant difference compared to Ver E-treated cells, as determined by two-way ANOVA with Sidak’s post hoc test.

## DISCUSSION

This study provides an initial biological and chemical characterization of Ver E, a marine-derived depsipeptide from *Okeania sp*^14^. Ver E exhibited several promising pharmacological properties: (1) modulation of calcium homeostasis without affecting SOCE, (2) direct, saturable binding to the σ₂R/TMEM97 with sub-nanomole percent affinity, (3) reduction of neuronal activity in hiPSC-derived nociceptors at physiological temperature, (4) no alteration of p-eIF2α expression in DRG neurons, and (5) no cytotoxic effects in HEK293 cells up to 30 µM. These findings support Ver E’s potential as a selective and non-toxic analgesic lead compound.

Our results position Ver E as a modulator of neuronal excitability with a distinct mechanism that sets it apart from other marine-derived compounds. The observed increase in intracellular Ca²⁺ levels is consistent with previous studies on cyanobacterial natural products^43^ but occurred independently of SOCE. This suggests that Ver E may act through non-canonical pathways, possibly involving direct modulation of plasma membrane ion channels or stimulation of Ca²⁺ release from intracellular stores unrelated to the STIM1-Orai1 SOCE system.

The transient suppression of spontaneous firing in hiPSC-derived nociceptors at physiological temperature, combined with a lack of effect during thermal stress, suggests that Ver E does not act on thermosensitive channels such as TRPV1. TRPV1 channels normally open in response to noxious heat to depolarize neurons^44–47^. Instead, Ver E’s selective inhibition of baseline excitability points to possible modulation of Naᵥ, Kᵥ, and Caᵥ channels, which are responsible for regulating resting membrane potential and the threshold for action potential generation. This level of selectivity is important in pain therapeutics, where broad inhibition of sensory pathways often results in undesirable side effects. Importantly, Ver E exhibited no cytotoxicity in HEK-293 cells, which is essential for establishing its safety profile and evaluating its potential as a therapeutic agent, particularly given that many marine-derived depsipeptides are known to exhibit neurotoxicity and cytotoxicity^48^. In summary, Ver E’s ability to modulate calcium dynamics in DRG neurons, its lack of effect on SOCE pathways, and its absence of cytotoxicity position it as a promising candidate for further investigation as a therapeutic agent targeting specific pain pathways.

### Ver E Modulates σ2R/TMEM97-dependent Cellular Responses

Ver E has low Maximum Tanimoto Scores (0.25 to 0.46, average of 0.32) when compared to previously described σ₂R ligands (**Table S1**). Tanimoto scores range from 0 to 1 with 1 being identical and 0 indicating no shared structural elements. Work by Patterson et al^49^ has been used to justify a Tanimoto value of 0.85 as highly likely to have similar or the same biological activity. Thus, the predicted and direct binding of Ver E to σ2R/TMEM97 coupled with the lower Tanimoto scores suggests that Ver E represents a structurally unique scaffold at this target. Ver E has a peculiar physicochemical profile, characterized by a relatively high predicted drug-likeness score, accompanied by unusually high molecular weight, topological polar surface area, and sp3 atom profile (**Table S2**). This combination of low Tanimoto score to other known ligands and high drug likeness score drive our enthusiasm for the veraguamide backbone as a scaffold. However, we need to understand its interactions with σ2R/TMEM97 better, so we evaluated it with computational docking and NMR binding experiments to begin to understand likely interactions driving the affinity.

Given our previous findings showing σ₂R/TMEM97 affinity for veraguamides^14^, we evaluated the potential for Ver E at this receptor. Both generative diffusion (**Figure S7**) and energy-based computational docking approaches (**Figure 2**) predicted binding in the entrance of the PB28^39^ and cholesterol binding pocket. While largely in the same region, the cyclic structure of Ver E precludes its binding in the pocket itself (**Figure 2E**, **Figure 2H**), providing an opportunity for side-chain modification to engage the internal pocket for specific and prolonged binding. Similar ‘anchor-extension’ remodeling of macrocycles has successfully deepened pocket engagement in other targets^50^. We also evaluated the binding affinity of Ver E to σ₂R/TMEM97, using a construct that lacks the ER retention tag, σ₂R-ΔER using solution NMR. This result is the first example of using NMR binding affinity on reconstituted human σ₂R/TMEM97. The results revealed specific, saturable binding with a global *K*_d_ of 0.3 ± 0.2 nmol%. The use of mol% accounts for the partitioning of Ver E between the DPC micelle environment and the bulk solution, providing a more accurate representation of the effective ligand concentration available for binding. The ^1^H-^15^N BEST-TROSY spectra (**Figure 3**) also showed broadening and disappearance of some resonances, suggesting conformational exchange. Upon titration with Ver E, selected resonances exhibited dose-dependent chemical shift perturbation (**S9**), confirming direct binding interaction with σ₂R/TMEM97. The control experiment (**S10**) demonstrated minimal solvent-induced effects, reinforcing the specificity of Ver E binding. Future studies will aim to identify whether Ver E binds to the σ₂R/TMEM97 orthosteric site. These first-in-class NMR binding data establish that Ver E binds specifically and with high affinity to σ₂R-ΔER, highlighting its promise as a lead compound for developing novel therapeutics targeting σ₂R/TMEM97 for pain management.

Although docking and NMR data indicate direct Ver E binding to σ₂R/TMEM97, our results using the ISR p-eIF2α assay complicate the story. In our previous publication, we demonstrated that FEM 1689 drives analgesic responses in mice with neuropathic pain through σ₂R/TMEM97 and that this response is associated with inhibition of the ISR that is absent in σ₂R/TMEM97 knockout mice^23^. In the present manuscript, we reproduced this inhibition effect of FEM 1689, but Ver E failed to modulate p-eIF2α. Given that Ver E shows potential to reduce activity in mouse DRG (**Figure 4**) and hiPSC-derived nociceptors (**Figure 5**), both of which would be expected to reduce pain, it is possible that there is another antinociceptive mechanism of σ2R/TMEM97 beyond ISR modulation. Alternatively, Ver E may have binding partners beyond σ2R/TMEM97.

### Limitations and future directions

While our study provides a comprehensive initial characterization of Ver E and suggests a favorable safety profile, several important questions remain. First, the precise molecular mechanisms by which Ver E modulates Ca²⁺ signaling have yet to be elucidated, underscoring the need for additional receptor-binding and electrophysiological studies to clarify its target specificity. Second, although we document Ver E’s *in vitro* efficacy, evaluation of the potential *in vivo* therapeutic benefits requires validation in animal models of chronic pain (e.g., inflammatory, cancer, and neuropathic). Evaluating its effects on thermal and mechanical sensitivity *in vivo* will further establish the clinical relevance of these findings.

Additionally, the transient nature of Ver E’s inhibitory effect on nociceptors suggests that structural modifications might be necessary to prolong its therapeutic duration, potentially guided by structure-activity relationship (SAR) studies. Finally, it remains critical to assess whether Ver E has off-target activities on other ion channels or signaling pathways to ensure a comprehensive safety profile. By addressing these limitations in future work, we can better define Ver E’s therapeutic potential and optimize it for clinical applications.

## Acknowledgments

This research was funded by the National Institutes of Health (NIH) grant NIH NINDS R61NS127271 (WVH, ECD, VK, BJK, KJT). This work was further supported by the Center for Pharmaceutical Research and Innovation (CPRI) as part of grant NIH P20 GM130456 (KJT).

## METHODS

### Cyanobacterial Collection and Isolation Sample Collection and Extraction

A cyanobacterial sample (DUQ0008) was collected from the coastal waters of Isla Mina in the Las Perlas Archipelago, Panama. A description of this collection with phylogenetic analysis of the collected species was previously published^14^. The sample containing veraguamide E (75 g, dry wt) was extracted using a CH₂Cl₂–MeOH (2:1) solvent mixture, yielding 3.3 g of crude extract.

### Chromatography

The crude extract underwent normal-phase chromatography on silica gel (230–400 mesh), using a stepwise gradient solvent system starting from 100% hexanes, then gradually increasing polarity by adding ethyl acetate, and finally adding methanol. This process yielded nine fractions (A–I). A portion of fraction G (15 mg), eluted with 70% methanol, was subsequently purified by reversed-phase HPLC (Phenomenex, Kinetex 5 μm, 150 × 10 mm) with a linear gradient of acetonitrile–0.1% TFA in water (60% acetonitrile for 5 min, 60–100% acetonitrile over 12 min, then 100% acetonitrile for 5 min), affording 2.8 mg of the target compound as a colorless oil (**S1, S2**). Semi-preparative HPLC separations were performed on an Agilent 1260 Infinity II system with UV detection using HPLC-grade solvents.

### Ver E NMR Spectroscopy

All NMR spectra were recorded in CDCl₃ (δ_C 77.2, δ_H 7.24) on a Bruker 600 (AV4 NEO) MHz spectrometer. The instrument was equipped with a 5 mm ^1^H-optimized triple-channel CryoProbe for ^1^H observation, with ^13^C/^15^N decoupling and ^13^C observation with ^1^H decoupling (**S5, S6**). The NMR data for the following compound has been deposited in the Natural Products Magnetic Resonance Database (NP-MRD; www.np-mrd.org) and can be found at NP0009879 (Ver E).

### Mass Spectrometry

High-resolution mass measurements were acquired on a SCIEX Q-TOF 5600 mass spectrometer with a dual-spray electrospray ionization source, connected to a Shimadzu DGU-20A 3R system. For MS/MS spectra, an IonSpray voltage floating 5500 V was used with a collision energy of 3500 V. The source temperature was maintained at 500 °C, with ion source gas 1 and 2 flows of 50 L/h and 55 L/h, respectively. MS/MS survey scans were run as a dependent acquisition, in which the 20 most intense ions in the MS spectrum were selected as MS/MS precursors. The threshold for MS/MS targeting was set at 100, with each targeted ion excluded from further targeting for 15 s.

### Molecular Networking

Molecular networks were generated using the online workflow at the GNPS2 website (http://gnps2.org). The data were clustered with a parent mass tolerance of 2 Da and an MS/MS fragment ion tolerance of 0.4 Da. Edges in the network were retained if they had a cosine score above 0.4 and more than six matched peaks. The resultant consensus MS/MS spectra were searched against GNPS2’s spectral libraries. A cosine similarity score of 0.4 was obtained when comparing the MS/MS fragmentation pattern of the purified compound with that of Ver E, resulting in 15 matching peaks (**S6**). In the dependent MS/MS acquisition, the 20 most intense ions in each MS survey scan were selected if they exceeded an intensity threshold of 100, after which they were excluded from reselection for 15 s. Data visualization for the molecular network was also conducted at the GNPS2 website. A summary of the spectral data is in the SI and publicly available on GNPS2 using the Task ID number 0075011f9ad94b779bb735618e11c278.

### Computational Analysis of Sigma-2 ligands and Ver E

Overlap Analysis to generate Maximum Tanimoto Scores was performed using an academic license of the Instant JChem platform (ver. 23.5.0, 2023). The overlap analysis with Tanimoto Score screening configuration was done using a similarity search mode based on a Chemical Hashed Fingerprint with a 0.01 threshold, where the Ver E was set as the *query* and selected σ2R ligands (**Table 1**) as the *target*. Predicted physicochemical values (**Table 2**) were determined using DataWarrior (ver. 06.04.02, 2025) and Principal Component Analysis was performed using RStudio (ver. 2024.12.1).

### *In Silico* Docking Sigma-2 ligands and Ver E

#### Ligand and Receptor Structure Representation

The Sigma-2 receptor sequence was obtained structure was prepared using AlphaFold3 on the AlphaFold server^38,51,52^. Two-dimensional representations of Veraguamides C, E, O, and P were rendered in three dimensions using the obgen tool^53^. The 3-dimensional representations were then assigned hydrogens and charges using “prepligand” python script for Dock6.11^54^.

#### Autodock-GPU Docking

To prepare the ligands for AutoDock-GPU, we converted the output .mol2 format to .pdbqt format using OpenBabel version 3.1.0 ^55.^ After the addition of hydrogens, the .pdbqt format of the receptor was converted to a .maps and .fld file format for docking using AutoGrid4, part of the AutoDockTools 1.5.7 release^56^. Ligand-receptor interactions were estimated using AutoDock-GPU, with 1000 runs / molecule specified^57,58^. Output scores were saved to .txt formats, which were read using a lab-developed Python script.

#### RosettaLigand Docking and Interaction Analysis

Conformers of the ligands screened were generated by using obconformer in a scripted loop, then concatenated into a single file using the “cat” unix shell function^55^. This conformer file was then processed into a Rosetta-readable .pdb file using the molfile_to_params Python script that is part of the Rosetta suite^59^. Docking was performed with parsed functions using Rosettascripts with the centerpoint based on the PB28 center, search radius of 20 Angstroms, and 10000 cycles^60,61^. The top-scoring pose was evaluated for interacting residues using the Protein-Ligand Interaction Profiler (PLIP) server^62^.

#### Diffusion Based Docking Analysis

Generative diffusion-based docking was performed using DiffDock^63,64^. We used the Nvidia NIM API for DiffDock (https://build.nvidia.com/mit/diffdock), part of the Nvidia BioNeMo framework^65^.

We generated 100 poses using the AlphaFold3 generated structure of Sigma2 for the Target Protein and the generated 3D representation of Veraguamide E as a ligand. All docking simulations were performed with 20 diffusion steps and 20 time divisions. Poses were scored by their confidence measure, with the top scoring pose evaluated for interacting residues using the Protein-Ligand Interaction Profiler (PLIP) server^62^.

### NMR Ligand Binding

#### Human σ_2_R Expression and Purification

The gene encoding the human σ_2_ receptor (σ2R) was cloned into the pETSG vector, which includes a C-terminal 8×His tag linked to thermostable GFP (TGP) followed by a thrombin cleavage site^66^. The σ2R gene was obtained from DNASU^67^ (plasmid # HsCD00514886) and pETSG was a gift from Prof. Dianfan Li (Addgene plasmid # 159418). The σ2R construct was truncated at the ER retention signal (residue Lys168, σ2R-ΔER) to improve expression. Whole plasmid sequencing (Plasmidsaurus) confirmed the pETSG-σ2R-ΔER identity.

σ₂R/TMEM97 expression was optimized for the BL21 (DE3) *E. coli* strain. A 5 mL LB overnight starter culture was prepared from a single colony, supplemented with 0.05 mg/mL kanamycin (Gold Biotechnology), and incubated at 37 °C. The starter culture inoculated 1 L of M9 minimal media (12.8 g Na_2_HPO_4_·7H_2_O, 3.0 g KH_2_PO_4_, 0.5 g NaCl, 1 g ^15^NH_4_Cl, 20 mL of 20% w/v D-glucose, 10 mL 100× MEM vitamin solution, 1 mL 1 mM MgSO_4_, 1 mL 0.1 mM CaCl_2_, 500 µL of 1000× trace metal mixture). The culture was grown at 25 °C, and protein expression was induced at OD_600nm_ of 0.5-0.6 with 1 mM isopropyl β-D-1-thiogalactopyranoside for 22 hours. The resulting cells were harvested by centrifugation at 6000 ×*g* (20 min, 4 °C).

The cell pellet was resuspended in 7 mL of lysis buffer (75 mM Tris-HCl (VWR), 300 mM NaCl (Fisher Scientific), 0.5 mM EDTA (Sigma Aldrich), pH 7.8) per gram of pellet supplemented with 0.1 mg/mL lysozyme, 0.01 mg/mL DNase, 0.01 m/mL RNase, 10 µL of 0.5 M magnesium acetate (Sigma Aldrich) and 10 µL of 100 mM PMSF (Sigma Aldrich) per mL of lysis buffer. The sample was tumbled at room temperature for 30 min and then sonicated at 4 °C (Qsonica S-4000 Ultrasonic Processor) with a 50% duty cycle (5 sec on/5 sec off) at 55% power. 3% v/v Empigen (N, N-Dimethyl-N-dodecylglycine betaine, BOC Sciences) was added to the cell lysate and tumbled at 4 °C for 1 hr to extract σ2R-ΔER into micelles. The lysate was then centrifuged (38,500 ×*g*, 25 min, 4 °C) to pellet cellular debris. The supernatant was incubated with 2 mL of preequilibrated Ni(II)-NTA Superflow (Qiagen) resin for 1–1.5 hr at 4 °C.

The σ₂R/TMEM97 bound resin was packed in a gravity column and washed with 20 column volumes of Buffer A (40 mM HEPES (Gold Biotechnology), 300 mM NaCl (Fisher Scientific), 2 mM β-ME (Sigma Aldrich), pH 7.8) with 1% (v/v) Empigen. The resin was washed with 20 column volumes of wash buffer (40 mM HEPES, 300 mM NaCl, 50 mM imidazole (Sigma Aldrich), 1% (v/v) Empigen, 2 mM β-ME, pH 7.8). Empigen was exchanged for n-dodecylphosphocholine (DPC, Anatrace) with at least 15 column volumes of rinse buffer (20 mM HEPES, 150 mM NaCl, 0.5% (w/v) DPC, 2 mM β-ME, pH 7.8). σ2R-ΔER was eluted with 10 column volumes of elution buffer (20 mM HEPES, 300 mM imidazole, 150 mM NaCl, 0.5% (w/v) DPC, 2 mM β-ME pH 7.8). The eluted protein from Ni^2+^-NTA column purification was buffer exchanged by ultrafiltration using a 10 kD cut-off Amicon Ultra-15 centrifugal filter (Millipore) with four 4 mL exchanges with buffer (20 mM HEPES, 150 mM NaCl, pH 7.8) followed by concentration step to 0.5 mL. Three units of thrombin (Novagen) were added to the sample, and the reaction was incubated for 22 hr at room temperature. Post-thrombin cleavage, the sample was passed over Ni-NTA resin, and the flow-through containing the thrombin-cleaved σ₂R/TMEM97 was collected. The cleaved protein was concentrated to 0.5 mL and further purified by gel filtration chromatography using a Superdex 200 Prep Grade Resin packed in a 16XK column preequilibrated with NMR buffer (20 mM HEPES, 150 mM NaCl, 0.5 mM EDTA, 0.2% (w/v) DPC, pH 6.8). SDS-PAGE was used to assess purity (**S9**), showing the combined fraction and concentrated to ∼175 µL for NMR studies. The average σ₂R/TMEM97 yield was ∼50 μg/L.

#### NMR-detected Veraguamide E Binding Studies

The NMR sample was prepared in a 3 mM NMR tube (Bruker) with a total volume of 180 µL, including 5 µL D_2_O. ^1^H-^15^N BEST-TROSY NMR experiments (b_trosyf3gpph.2 pulse program)^68^ were recorded on a Bruker Avance III HD 850 MHz spectrometer equipped with a cryogenically cooled 5 mm TCI cryoprobe. Titrations were diluted from 1 mM stock Ver E prepared in 200 proof ethanol (VWR), and concentrations were measured in mol% and calculated using the equation:

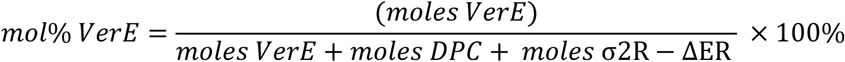

For NMR experiments, a sample containing 53 μM σ₂R/TMEM97 in 4.5% DPC (w/v) was used, and a series of ^1^H-^15^N BEST-TROSY spectra were recorded as a function of Ver E at 37 °C, with 88 scans per spectrum. The Ver E concentrations used were 0.003, 0.03, 0.3, 0.6, 3, and 7.5 nmol%. NMR data were processed in nmrPipe^69^ and analyzed with the CcpNMR Analysis software^70^. Chemical shift perturbations induced by Ver E were calculated by:

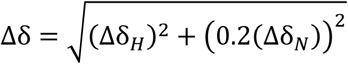

where Δδ_*HH*_ and Δδ_*NN*_ are the proton and nitrogen chemical shift position differences between the initial titration point (0 nmol% Ver E) and a specific Ver E concentration for a given resonance^71^. For selected resonances, when Δδ values were plotted as a function of Ver E concentration, saturable binding isotherms were observed, indicating specific binding of Ver E to σ₂R/TMEM97. TREND NMR^72,73^ was applied, and PCA was used to analyze titration data globally, where PC1 captures the greatest variance and serves as an indicator of ligand binding^74^. The dissociation constant, *K*_d,_ was calculated by fitting (SigmaPlot) the PCA data to a single-site binding model^71,75^ :

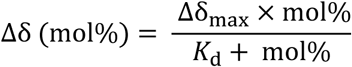

where Δδ*_max_* is the maximum chemical shift perturbation.

A control binding experiment with the vehicle ethanol (EtOH) was performed following a parallel procedure as above to rule out that EtOH binds to σ₂R/TMEM97 (**S10**).

#### Animals and Ethics

All animals used for the Ca²⁺ imaging experiments were housed in groups of three to four per cage, maintained under a 12-hour light/dark cycle, and provided food and water *ad libitum*. All protocols and procedures were conducted in compliance with protocol 20-04, approved by the Institutional Animal Care and Use Committee (IACUC) at the University of Texas at Dallas.

#### Calcium Imaging Experiments

An 18-week-old male mouse, heterozygous for GCamp6f and positive for Nes-Cre recombinase, was used for the wild-type fluorescent experiments. To generate Nestin-GCaMP6f transgenic mice for Ca²⁺ imaging, C57Bl/6J mice (Jackson Laboratories, Bar Harbor, ME) expressing a Cre-dependent GCaMP6f calcium sensor (B6J.Cg-Gt(ROSA)26Sortm95.1(CAG-GCaMP6f)Hze/MwarJ, JAX #028865) were crossed with C57Bl/6J mice carrying a Nestin promoter-driven Cre recombinase (B6.Cg-Tg(Nes-cre)1Kln/J, JAX #003771). Offspring were then genotyped using primers from Integrated DNA Technologies, Inc.

#### Isolation of Primary Mouse Sensory Neurons

Primary dorsal root ganglion (DRG) neurons were prepared and cultured as follows. Twelve-millimeter glass coverslips (Bellco Glass, cat# 1943-10012A) were spot-coated with 70 µL of 70 µg/mL poly-D-lysine (PDL; Sigma-Aldrich, cat# P7405) and incubated overnight at 37 °C with 5% O₂. After incubation, coverslips were rinsed three times with sterile water (Sigma-Aldrich, cat# 95284) and dried in a laminar flow hood at room temperature. Next, a mouse was decapitated, and its spinal column was removed and placed in ice-cold Hanks’ Balanced Salt Solution (HBSS; Fisher Scientific, cat# 14-170-161) supplemented with 10 mM HEPES (Sigma-Aldrich, cat# H3375). The spinal cord was bisected, and each half was transferred to ice-cold Hibernate A solution (Fisher Scientific, cat# A1247501). All DRGs were extracted into a 5 mL snap-cap tube containing ice-cold Hibernate A.

To enzymatically dissociate the DRGs, papain was prepared by mixing 2 mL of an “HBSS complete” solution (48.5 mL of HBSS plus 0.5 mL penicillin/streptomycin, 0.5 mL sodium pyruvate [Fisher Scientific, cat# 11-360-070], and 0.5 mL HEPES) with papain (Worthington Biochemical, cat# LS003119) to achieve a final concentration of 20 IU/mL. The papain solution was filtered (0.20 µm) and kept at 37 °C until use. The DRGs were treated with 15 IU/mL of this papain mixture plus 2.25 mg/mL collagenase (Sigma-Aldrich, cat# C6885) at 37 °C for 20 minutes. The resulting cell suspension was filtered through a 40 µm nylon mesh (Fisher Scientific, cat# 08-771-1), centrifuged at 160 rcf for 4 minutes, washed once, and pelleted. The pellet was resuspended in Neurobasal A medium (Thermo Scientific, cat# 10888022) supplemented with 5% HI fetal bovine serum (VWR, cat# 10802-772), B-27 (Thermo Scientific, cat# A3582801), 2 mM GlutaMax-1 (Thermo Scientific, cat# 35050061), and 50 IU/mL penicillin/streptomycin (Thermo Scientific, cat# 15070063). Cells were counted (1:1 in trypan blue) and plated at 450,000–500,000 cells/mL. After four hours of attachment at 37 °C with 5% O₂, 400 µL of warm, complete Neurobasal A medium was added to each coverslip, and the plates were incubated overnight.

#### Fluorescent Imaging of Mouse Sensory Neurons

Calcium imaging was performed 24 hours post-incubation using an upright Olympus BX51WI microscope equipped with a 480 nm interface filter, a 505 nm dichroic mirror, and a 535 nm barrier filter (FITC band-pass). Images were collected at 1 Hz with Cell-Sense software (Olympus) and an Orca Fusion C14440 sCMOS camera (Hamamatsu). Nociceptive cells were identified in the Cell-Sense software, yielding over 100 distinct regions of interest (ROIs). Changes in intracellular Ca²⁺ were tracked over time.

To apply compounds or vehicle controls, a switching valve system (VC-6, Warner Instruments) was employed for bath perfusion into a 358 µL recording chamber (Warner Instruments, RC-21B), regulated by nitrogen gas (VPP-6, Warner Instruments). Coverslips were initially bathed in a baseline recording solution, followed by test compound or vehicle application, and then rinsed again with the baseline solution. At the conclusion of each recording, 60 mM potassium chloride (KCl) was introduced as a positive control for neuronal activation. Neurons responsive to KCl were statistically analyzed across coverslips, focusing on the peak Ca²⁺ signal and area under the curve (AUC) for each treatment group.

#### Screening for the Analgesic Properties of Ver E in hiPSCs

To evaluate the analgesic properties of Ver E, a multi-well microelectrode array (MEA) system was employed using hiPSC-derived nociceptors^28^. Prior to cell seeding, a 48-well MEA plate (Axion Biosystems, cat# M768-tMEA-48W) underwent a standardized surface-modification protocol: spot-coating the wells with 0.01% Poly-L-Ornithine (EMD Millipore Sigma, cat# A-004-C) and incubating overnight at room temperature. The plate was then rinsed three times with sterile deionized water. A biocompatible coating—iMatrix-511 silk (Iwai-chem, cat# SKU:N-892022)—was diluted 1:50 in DPBS (Sigma-Aldrich, cat# D8537) and spot-applied to each well, followed by a three-hour incubation at 37 °C.

Chrono™ Sensory Neurons (Anatomic™, cat# 7009) were thawed, washed in DMEM/F12 (gibco™, cat# 11320-033), and resuspended in Chrono™ Senso-MM complete growth medium (Anatomic, cat# 1030). Excess iMatrix-511 was removed, and the cells were spot-seeded (5 µL/well). After a 20-minute attachment period, Chrono™ Senso-MM medium was gently added to each well (final volume: 400 µL). The medium was partially exchanged (50%) the following day and subsequently every 2–3 days. Cultures were maintained for four weeks at 37 °C in a humidified incubator with 5% CO₂.

Electrophysiological recordings were performed on an Axion Maestro Pro system at a 12.5 kHz sampling rate. The data were processed with a single-pole Butterworth bandpass filter (300–5000 Hz) to isolate neuronal signals. Spikes were defined as voltage deflections exceeding ±5.5 SD of the baseline root mean square (RMS), with a minimum firing rate set at 1 spike per minute. At DIV28, baseline recordings were taken at both 37 °C and 42 °C, followed by administration of Ver E at 1–30 µM. Activity was measured for 15 minutes post-treatment, then at 1 hour, 2 hours, and 24 hours, with all time points (except 0–0.25 hours) recorded at both temperatures. After 24 hours, wells were washed, and a final measurement was taken at least 1 hour later at 37 °C. For each time point, total spike counts per well were normalized to their respective baselines. Statistical analysis used one-way ANOVA with Dunnett’s post hoc test (p < 0.05 considered significant).

#### Immunocytochemistry and p-eIF2α Detection in Primary Mouse DRGs

Primary mouse DRGs were extracted and plated as described above, then maintained overnight in a complete Neurobasal A medium. After approximately 16 hours of treatment with Ver E, FEM 1689, or other small molecules, the culture media were removed, and the DRG neurons were fixed by adding 50 µL of 10% formalin in PBS for 10 minutes. The cells were washed twice with 1× PBS before a blocking/permeabilization step, which involved a 1-hour incubation at room temperature with 50 µL of 10% normal goat serum (NGS) in 0.1% Triton X–PBS. Following one additional PBS wash, primary antibodies were introduced in a solution containing 0.1% PBS– Triton X, NGS, and bovine serum albumin.

Specifically, a rabbit anti–p–eIF2α (Cell Signaling, cat# 3398) was used at 1:500, and a chicken anti-peripherin (Encor Biotechnology, cat# CPCA-Peri) was used at 1:1000 for primary immunodetection. After overnight incubation at 4 °C, coverslips were washed twice with 0.5% Tween in PBS and once with PBS alone. Cells were then incubated for 1 hour at room temperature with secondary antibodies diluted in the same buffer. Secondaries included a goat anti-rabbit Alexa Fluor 555 (Thermo, cat# A-21428) at 1:1000 and a goat anti-chicken IgY Alexa Fluor 488 (Thermo, cat# A11039) at 1:2000. After one wash in 0.5% Tween in PBS and two washes in PBS, coverslips were inverted onto microscope slides containing 10 µL of antifade hard-mount DAPI (Fisher, cat# NC9029229) to stain nuclei. Coverslips were stored overnight at 4 °C in the dark. Fluorescence was visualized using an Olympus IX73 microscope with the following filter sets: TRITC (540–570 nm) for p-eIF2α, UV (350–405 nm) for DAPI, and GFP (480– 500 nm) for peripherin. Neurons were identified by peripherin staining, and p-eIF2α levels were quantified by measuring the mean gray intensity in regions of interest.

#### Characterization of the Cytotoxic Profile of Ver E

HEK-293 cells^76^ (generously provided by Dr. Xintong Dong, University of Texas at Dallas) were cultured in Dulbecco’s Modified Eagle Medium (DMEM; Cytiva, cat# SH30023.FS) supplemented with 10% fetal bovine serum (VWR, cat# 10802-772) and 100 U/mL penicillin-streptomycin (Gibco, cat# 15-070-063). The cells were maintained at 37 °C in a humidified atmosphere containing 5% CO₂ until they reached approximately 90% confluence. For experimental assays, cells were dissociated with 0.05% trypsin (Fisher Scientific, cat# 15-400-054) for 5 minutes, then neutralized with fresh culture media. The suspension was centrifuged at 300g for 5 minutes, after which the pellet was resuspended in new culture media.

In 96-well plates, HEK-293 cells were seeded at a density of 6 × 10^4 cells per well and allowed to adhere for 24 hours. Thereafter, cells were treated with Ver E or digitonin, each dissolved in DMEM, at final concentrations ranging from 0.03 µM to 30 µM. Vehicle-only controls received 0.25% DMSO (VWR, cat# BDH1115-1LP). Following 24 hours of compound exposure, cell viability was evaluated using a resazurin reduction assay, wherein living cells convert resazurin to the fluorescent compound resorufin. Resazurin sodium salt (Sigma Aldrich, cat# 62758-13-8) was freshly prepared at 0.15 mg/mL in DPBS (pH 7.4; Sigma-Aldrich, cat# D8537), filter-sterilized (0.2 µm), and protected from light. A 20 µL aliquot of this resazurin solution was added to each well, containing cells in 100 µL of media. After a 4-hour incubation at 37 °C, fluorescence was measured at an excitation wavelength of 560 nm and an emission wavelength of 590 nm on a microplate reader. Cell viability was determined by comparing the fluorescence of compound-treated wells to that of vehicle-treated controls.

### Statistics

#### *In Vitro* Calcium Imaging

All statistical analyses were performed using GraphPad Prism (version 9.0) and MATLAB. Raw data files, generated after each compound application by Cell-Sense software (Olympus), were transferred into Microsoft Excel and processed via a custom MATLAB script. This script calculated changes in fluorescence over time (ΔF/F₀), the peak fluorescence during compound application (max ΔF/F₀), and the area under the curve (AUC). The first region of interest (ROI), which contained an empty background, was used for subtraction in all other ROIs. The MATLAB script accounted for specific drug application time windows, and the resulting processed data were compiled into new spreadsheets for import into GraphPad Prism, where final visualizations and statistical tests were conducted. In the Ver E assays, unpaired two-tailed t-tests were used to compare 10 µM Ver E with 0.1% DMSO vehicle versus vehicle alone in two distinct parts of the recording (direct Ver E and SOCE response).

#### Screening for the Analgesic Property of Ver E

At each experimental time point, the total spike count in each well was normalized to its respective baseline (37 °C or 42 °C). Statistical comparisons were performed using a two-way ANOVA with Dunnett’s post hoc test to evaluate different Ver E concentrations (1–30 µM) against DMSO vehicle controls. Data are reported as mean ± SEM, with a p-value < 0.05 considered statistically significant. At 37 °C, Ver E at 30 µM induced a notable reduction in firing activity (p < 0.05), whereas no significant effect was observed at 42 °C.

#### Immunocytochemistry and p-eIF2α Detection in Primary Mouse DRGs

Mean gray intensities corresponding to p-eIF2α staining were calculated for each treatment group and are expressed as mean ± SEM. One-way ANOVA followed by Tukey’s multiple-comparisons test was used to assess statistical differences among Ver E (100 nM, 1000 nM), FEM 1689 (30 nM, 100 nM), and vehicle.

#### Characterization of the Cytotoxic Profile of Ver E

A logarithmic non-linear regression curve *[log⁡(inhibitor concentration) vs. normalized response]* was generated in GraphPad Prism 10 to determine the LC₅₀ of both Ver E and digitonin. The IC₅₀ calculated by this curve-fitting method was taken to represent the LC₅₀ in this experiment. Digitonin yielded an LC₅₀ value of 9.68 µM, whereas Ver E’s LC₅₀ exceeded 30 µM. No additional statistical comparison was required to confirm that Ver E did not induce detectable cytotoxicity within the 0.03–30 µM range.

