## Supplemental Figures for "Neurobiological and Chemical Characterization of the Cyanobacterial Metabolite Veraguamide E"

### Supplemental Figures and Tables

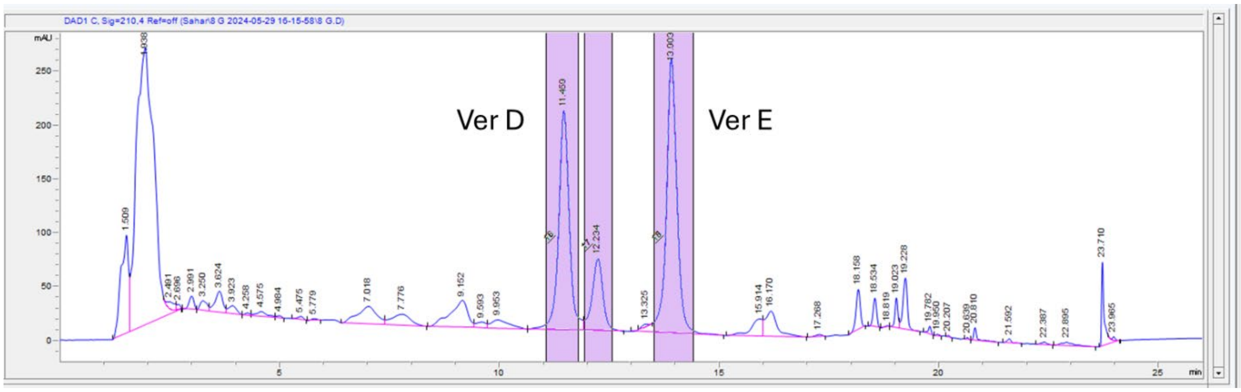


**Figure S1.** Preparative HPLC spectrum of DUQ0008-G for isolation of Ver E

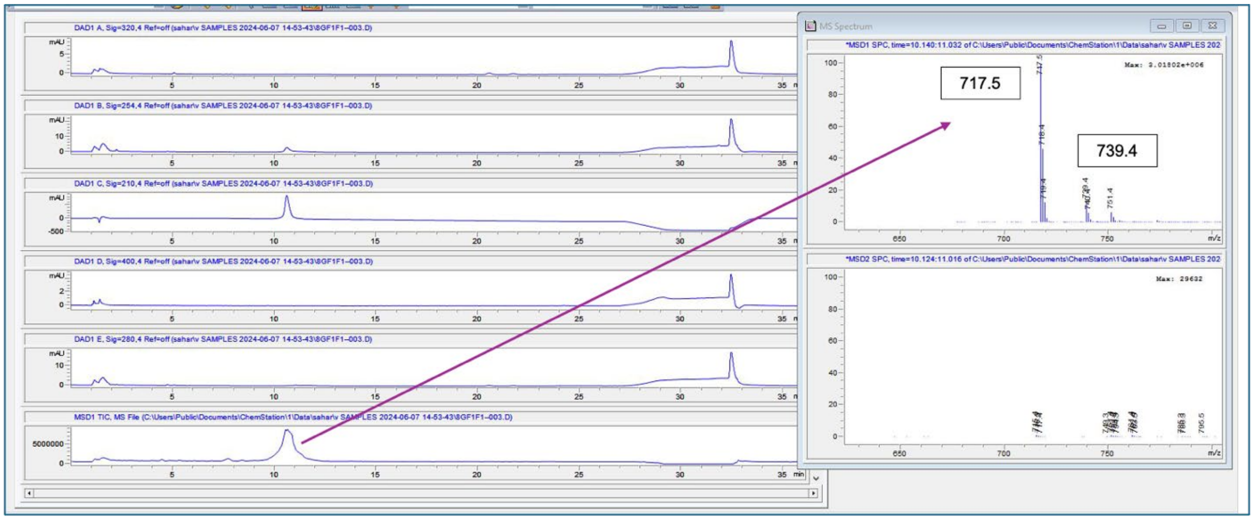


**Figure S2.** LC-MS spectrum of purified VerE

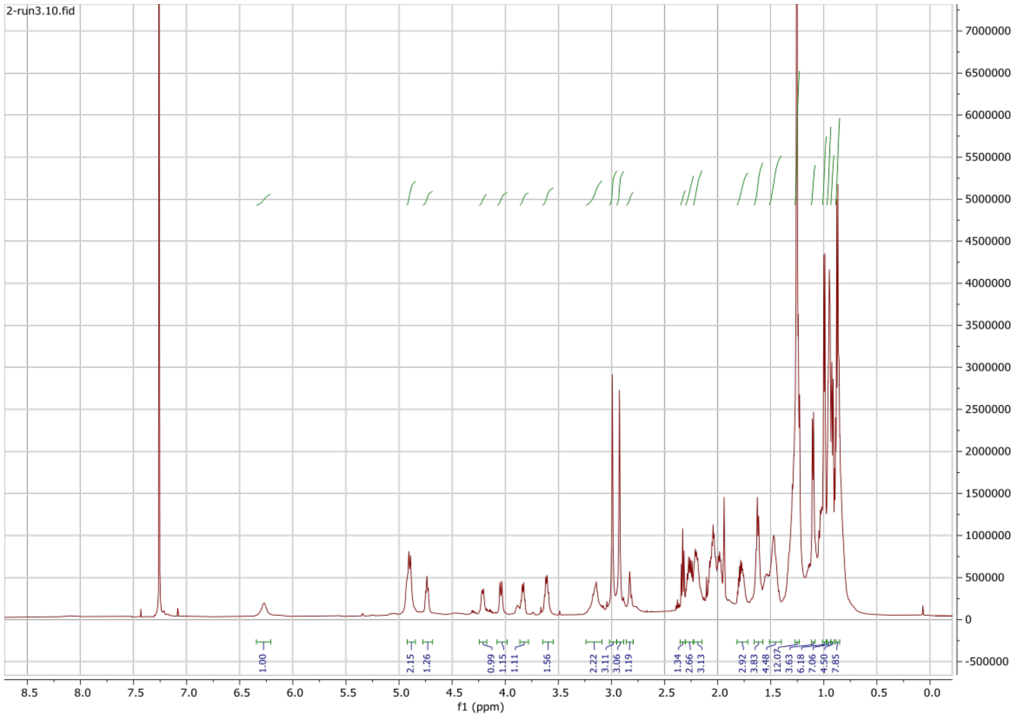


**Figure S3.** 1H NMR spectrum of purified Ver E in $CDCl_3$

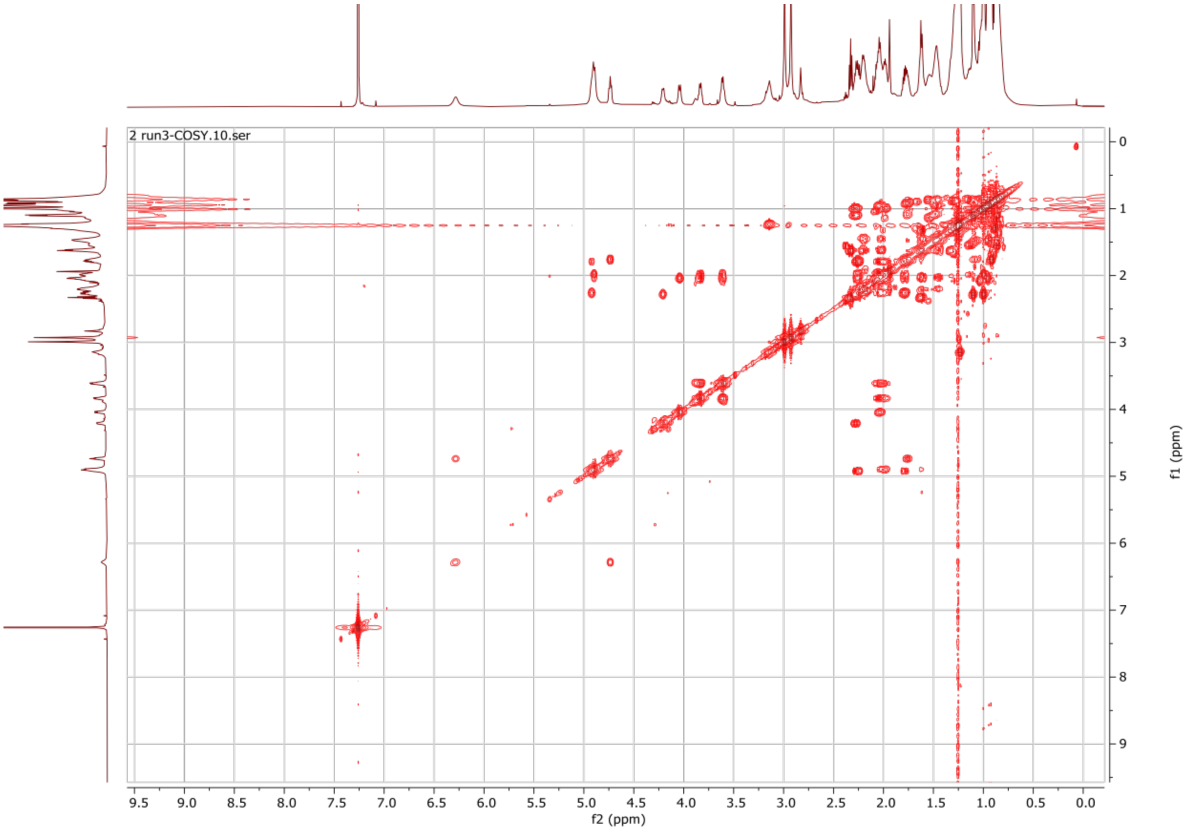


**Figure S4.** COSY spectrum of purified Ver E in $CDCl_3$

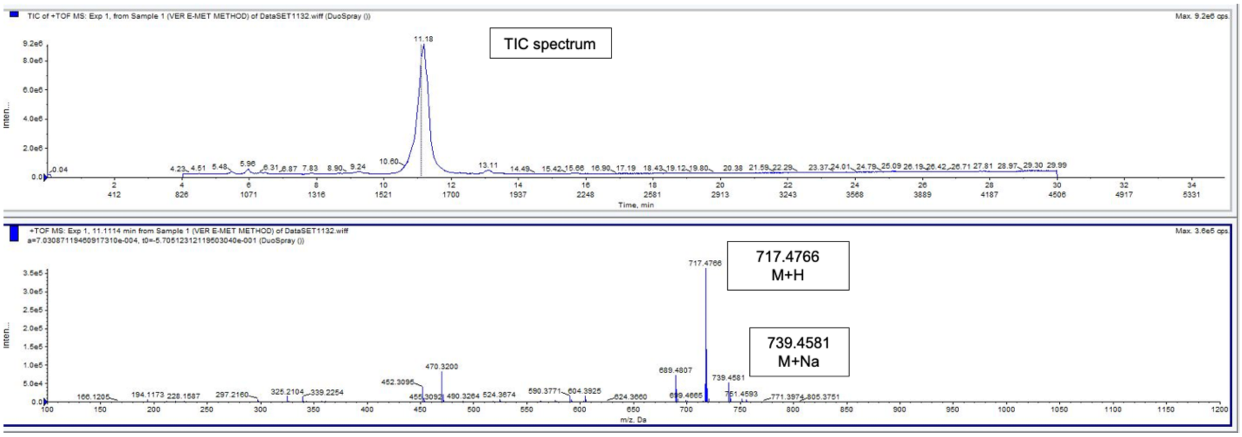


**Figure S5.** HRMS spectrum of purified ver E confirming the M+H (717.4766) and M+Na (739.4581)

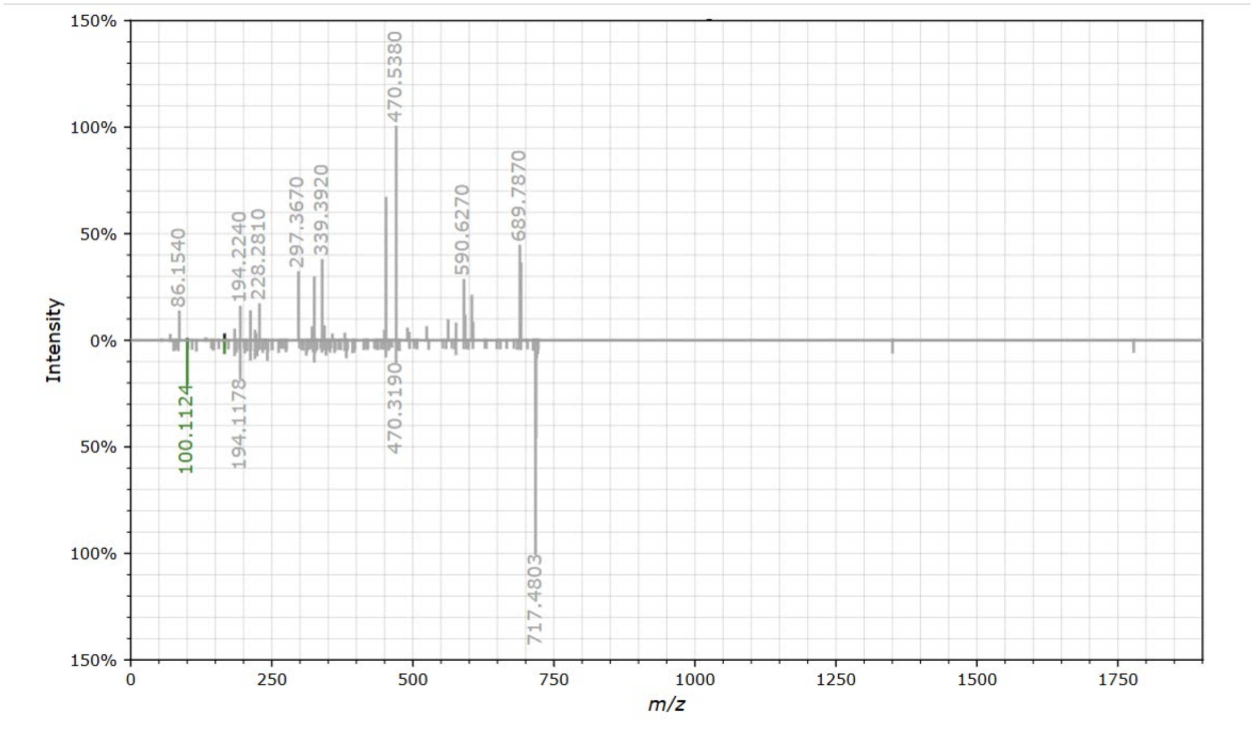


**Figure S6.** MS/MS spectral comparison (experimental (up) vs. library (down)) for Ver E.

**Table S1. Maximum Tanimoto Scores for Select $\sigma_2$R/TMEM97 Ligands Compared to Ver E**

| $\sigma_2$R Ligand | Tanimoto Score (Overlap with Ver E) |
|---|---|
| Siramesine (Lu 28-179) | 0.34 |
| Ibogaine | 0.28 |
| CB-64D | 0.30 |
| SM21 | 0.41 |
| WC26 | 0.35 |
| SW120 | 0.37 |
| RHM-4 | 0.31 |
| CM398 | 0.29 |
| AMA-1127 | 0.31 |
| DKR-1677 | 0.32 |
| JVW-1034 | 0.25 |
| FEM-1689 | 0.29 |
| Z4857158944 | 0.31 |
| Z1665845742 | 0.31 |
| PB28 | 0.31 |
| CT1812 (Elayta) | 0.26 |
| Histatin-1 | 0.46 |
| **Median** | **0.31** |
| **Max.** | **0.46** |
| **Min.** | **0.25** |

#### Table S2. Predicted Physicochemical Properties of Select $\sigma_2$R/TMEM97 Ligands and VerE

| Entry | MW[a] (g/mol) | cLogP[b] | HA | HD | PSA[c] (Å$^2$) | Drug-likeness | sp3-Atoms | Shape Index | cLogS | Relative PSA | Stereo-centers | Aro-matic Rings | Electro neg-ative Atoms |
|---|---|---|---|---|---|---|---|---|---|---|---|---|---|
| JVW-1034 | 311.855 | 5.3946 | 1 | 0 | 3.24 | 2.256 | 9 | 0.54545 | -4.445 | 0.014835 | 2 | 2 | 2 |
| PB28 | 370.579 | 5.2618 | 3 | 0 | 15.71 | 2.5326 | 21 | 0.62963 | -3.9 | 0.055697 | 1 | 1 | 3 |
| Siramesine (Lu 28-179) | 454.587 | 5.8081 | 3 | 0 | 17.4 | 2.9887 | 12 | 0.58824 | -7.962 | 0.058263 | 0 | 4 | 4 |
| FEM-1689 | 361.406 | 5.0399 | 2 | 1 | 23.47 | -6.1601 | 11 | 0.57692 | -4.931 | 0.064115 | 2 | 2 | 5 |
| Ibogaine | 310.439 | 4.0357 | 3 | 1 | 28.26 | 3.9995 | 14 | 0.52174 | -3.966 | 0.11754 | 4 | 2 | 3 |
| AMA-1127 | 423.53 | 4.4976 | 5 | 0 | 36.02 | 0.58082 | 14 | 0.58065 | -4.113 | 0.10574 | 2 | 2 | 6 |
| DKR-1677 | 419.567 | 4.9155 | 5 | 0 | 36.02 | 1.8129 | 15 | 0.6129 | -4.099 | 0.10333 | 2 | 2 | 5 |
| SM21 | 337.846 | 3.4245 | 4 | 0 | 38.77 | 3.6871 | 14 | 0.6087 | -3.95 | 0.1446 | 4 | 1 | 5 |
| CB-64D | 333.43 | 3.5804 | 3 | 1 | 40.54 | 5.5698 | 9 | 0.52 | -3.769 | 0.11628 | 2 | 2 | 3 |
| CM398 | 395.501 | 3.2504 | 6 | 0 | 45.25 | 1.2211 | 13 | 0.58621 | -4.181 | 0.14196 | 0 | 2 | 6 |
| Z166584574 | 347.461 | 3.0338 | 4 | 2 | 52.15 | 3.6197 | 9 | 0.65385 | -3.643 | 0.14921 | 1 | 3 | 4 |
| Z485715894 | 338.449 | 2.3436 | 4 | 2 | 52.57 | 4.6664 | 10 | 0.6 | -3.503 | 0.1545 | 1 | 2 | 4 |
| WC26 | 437.582 | 4.3542 | 6 | 1 | 54.04 | -7.9785 | 16 | 0.5625 | -5.293 | 0.1496 | 3 | 2 | 6 |
| CT1812 (Elayta) | 431.595 | 3.4278 | 5 | 1 | 75.22 | -10.797 | 16 | 0.53333 | -4.485 | 0.17037 | 0 | 2 | 6 |
| RHM-4 | 570.418 | 3.1171 | 8 | 1 | 78.49 | 0.40379 | 17 | 0.57576 | -5.282 | 0.20503 | 0 | 2 | 9 |
| Veraguamide E | 716.957 | 4.4668 | 12 | 1 | 142.63 | 4.3387 | 33 | 0.35294 | -5.066 | 0.20917 | 10 | 0 | 12 |
| SW120 | 622.764 | 6.846 | 11 | 2 | 147.57 | -15.215 | 24 | 0.62222 | -7.485 | 0.25664 | 3 | 3 | 12 |
| Histatin-1 | 1428.43 | -6.1883 | 37 | 21 | 619.33 | -7.2533 | 33 | 0.42157 | -4.249 | 0.44392 | 9 | 4 | 37 |

[a]MW = molecular weight; [b]cLogP = calculated logarithm of the partition coefficient; [c]PSA = polar surface area; H-A = hydrogen bond acceptors; H-D = hydrogen bond donors; cLogS = calculated logarithm of solubility

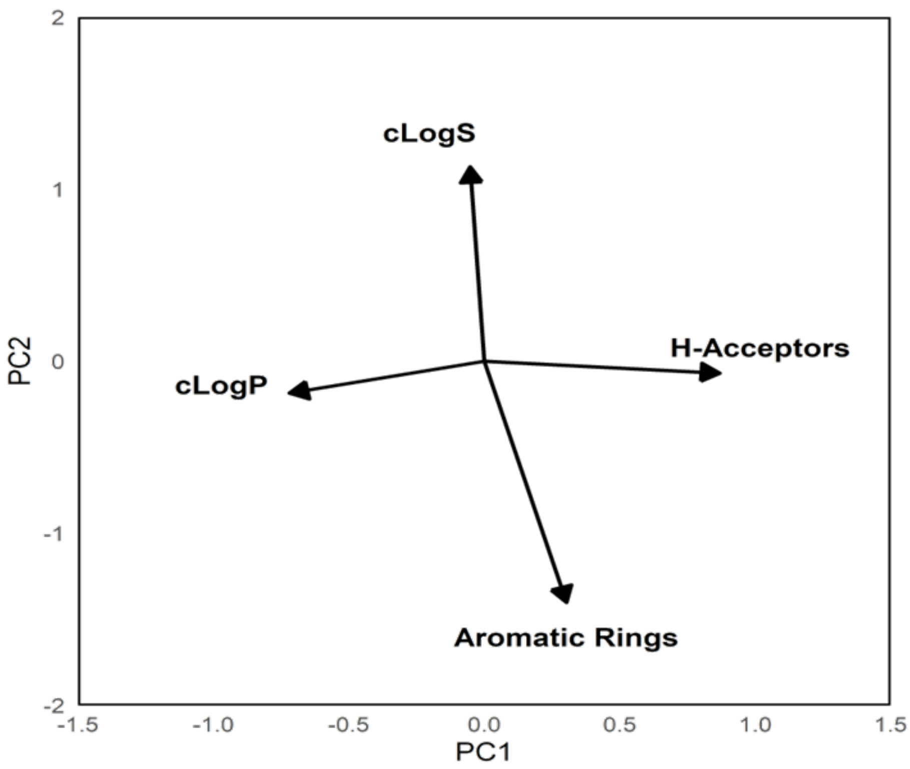


**Figure S7. PCA Loading Plot of 13 predicted physicochemical descriptors of VerE and previously described $\sigma_2$R ligands.**

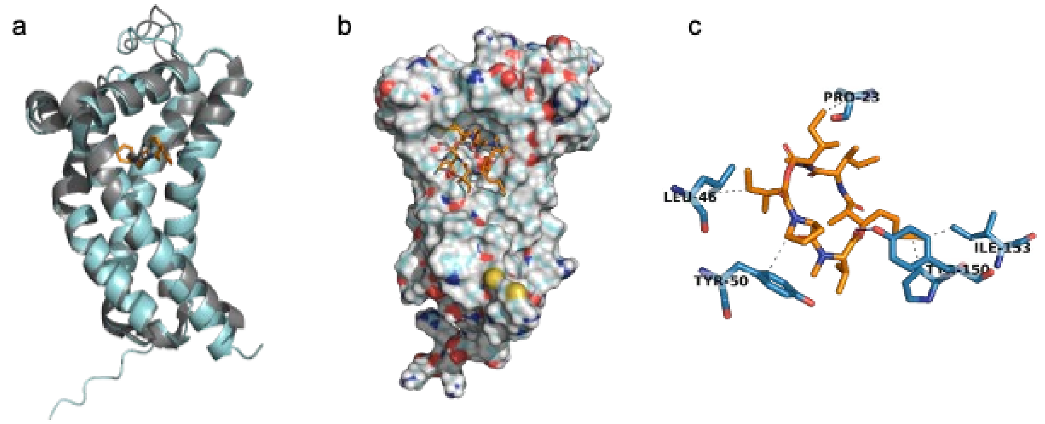


**Figure S8. Diffusion Model Benchmarking and Prediction. a)** DiffDock predicted (orange) vs. resolved (gray) structures of PB28 against bovine (resolved, gray) or Alphafold3 Predicted (teal) $\sigma_2$R/TMEM97. **b)** Predicted Ver E binding pose with **c)** key residues highlighted.

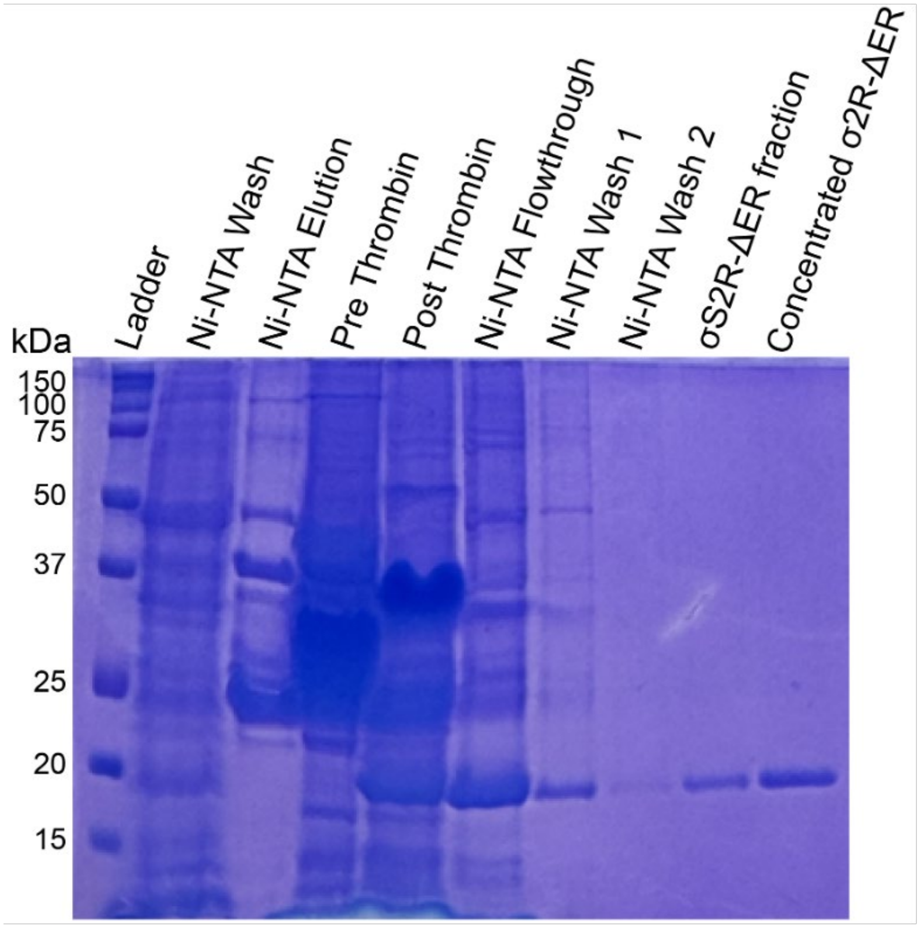


**Figure S9. SDS-PAGE gel of purified $\sigma_2$R/TMEM97 following Ni-NTA affinity chromatography and gel filtration purification.** The Ni-NTA elution fraction shows human $\sigma_2$R/TMEM97 (σ2R-ΔER) linked to the TGP protein (48 kDa). Following successful thrombin cleavage, the post thrombin fraction shows the anticipated shift in molecular weight, with the cleaved $\sigma_2$R/TMEM97 (~ 21 kDa) band appearing at the expected molecular weight. The Ni-NTA flowthrough and Wash 1 fractions contain the cleaved σ2R-ΔER, as indicated by the presence of the corresponding protein bands. Gel filtration further purified the protein, resulting in the isolated pure σ2R-ΔER for NMR studies.

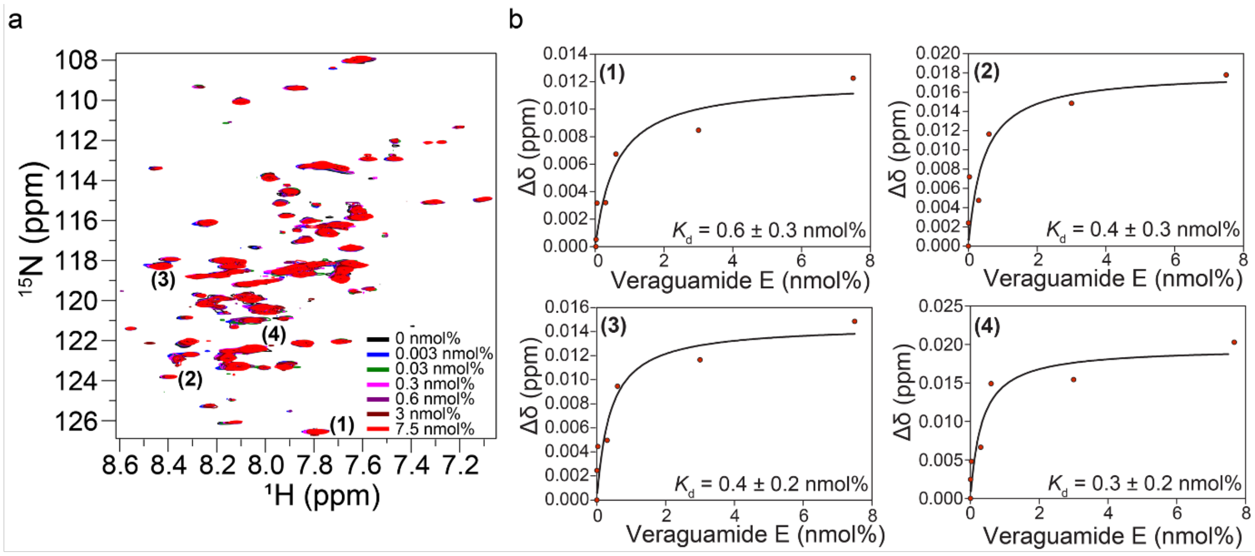


**Figure S10. Chemical shift perturbation analysis of selected resonances upon veraguamide E binding. a)** $^{1}H$-$^{15}N$ BEST-TROSY spectra of $\sigma_2$R/TMEM97 (σ2R-ΔER) titration with veraguamide E at 37 °C, with selected resonances highlighted to illustrate chemical shift perturbations. **b)** Selected resonances (1-4) show saturable binding isotherms as a function of veraguamide E.

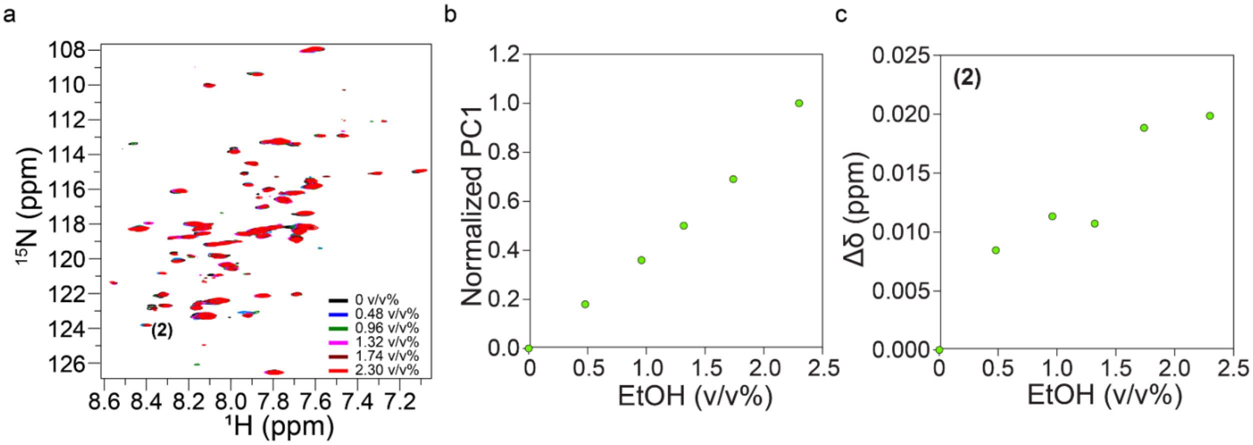


**S 11. The vehicle ethanol does not bind $\sigma_2$R/TMEM97 specifically. a)** Superimposed $^{1}H$-$^{15}N$ BEST-TROSY spectra of $\sigma_2$R/TMEM97 (σ2R-ΔER) titration with ethanol at 37 °C. **b)** Global analysis of chemical shift changes in response to EtOH concentrations, confirms that EtOH does not induce binding-related perturbations. **c)** Chemical shift perturbation analysis of a selected resonance further demonstrates that EtOH does not bind specifically to σ2R-ΔER.
